# A multiscale computational study of the conformation of the full-length intrinsically disordered protein MeCP2

**DOI:** 10.1101/2021.11.08.467619

**Authors:** Cecilia Chávez-García, Jérôme Hénin, Mikko Karttunen

## Abstract

The malfunction of the Methyl CpG binding protein 2 (MeCP2) is associated to the Rett syndrome, one of the most common causes of cognitive impairment in females. MeCP2 is an intrinsically disordered protein (IDP), making its experimental characterization a challenge. There is currently no structure available for the full-length MeCP2 in any of the databases, and only the structure of its MBD domain has been solved. We used this structure to build a full-length model of MeCP2 by completing the rest of the protein via *ab initio* modelling. Using a combination of all-atom and coarse-grained simulations, we characterized its structure and dynamics as well as the conformational space sampled by the ID and TRD domains in the absence of the rest of the protein. The present work is the first computational study of the full-length protein. Two main conformations were sampled in the coarse-grained simulations: a globular structure similar to the one observed in the all-atom force field and a two-globule conformation. Our all-atom model is in good agreement with the available experimental data, predicting amino acid W104 to be buried, amino acids R111 and R133 to be solvent accessible, and having 4.1% of α-helix content, compared to the 4% found experimentally. Finally, we compared the model predicted by AlphaFold to our Modeller model. The model was not stable in water and underwent further folding. Together, these simulations provide a detailed (if perhaps incomplete) conformational ensemble of the full-length MeCP2, which is compatible with experimental data and can be the basis of further studies, e.g., on mutants of the protein or its interactions with its biological partners.

## INTRODUCTION

Methyl CpG binding protein 2 (MeCP2) is a transcriptional regulator essential for growth and synaptic activity of neurons^1^. The malfunction of this protein is associated to the Rett syndrome, one of the most common causes of cognitive impairment in females^2,3^. The MeCP2 gene is X-linked in mammals. Mutations that affect the protein function were initially thought to be lethal in males^4^, but these are now frequently identified in cognitively impaired male patients^5^.

MeCP2 is an intrinsically disordered protein (IDP), and little is known about its molecular architecture during normal cellular processes and in disease^6^. IDPs are characterized by a low proportion of bulky hydrophobic amino acids and high proportions of charged and hydrophilic amino acids. Consequently, they cannot bury sufficient hydrophobic core to fold spontaneously into stable, highly organized three-dimensional structures; instead, they fluctuate through an ensemble of conformations^7^. The physical characteristics of IDPs makes their structural characterization a challenge as these proteins are more sensitive to degradation.

MeCP2 contains 486 amino acids, is a monomer in solution and is composed of six different domains^8^. Residues 78-162 specifically bind to methylated CpG dinucleotides and have been termed the methyl-CpG binding domain (MBD)^9^. Another functionally annotated region corresponds to the transcriptional repression domain (TRD) whose main function is to repress the transcription of genes^10^. Biophysical and protease digestion experiments identified three other domains: the N-terminal domain (NTD), the intervening domain (ID) and the C-terminal domain (CTD), which can be subdivided into CTD-α and CTD-β8 (Fig. 1).

**Figure 1.**
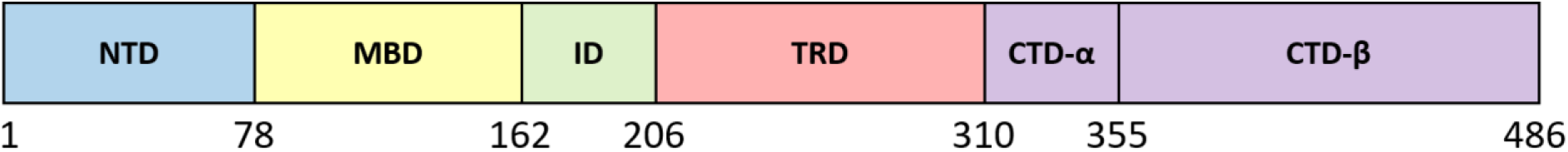
MeCP2 is composed of six domains: The N-terminal domain (NTD), the methyl-CpG binding domain (MBD), the intervening domain (ID), the transcription repression domain (TRD) and the C-terminal domain (CTD) which can be subdivided into CTD-α and CTD-β. The only available structure^1^ contains solely the MBD domain, which is the only ordered region in the protein.

There is currently no structure available for the full-length MeCP2 in any of the protein databases. MBD is the only domain for which the secondary structure is known, and it only accounts for ~17% of the amino acids^1^; MBD is also the only ordered domain. Circular dichroism (CD) of recombinant human MeCP2 has shown that the protein consists of ~35% β-strand/turn, 5% α-helix and almost 60% is unstructured^2^. Characterization of MeCP2 by hydrogen/deuterium exchange has indicated disorder in the entire polypeptide chain with the exception of the MBD domain^3^. Further CD studies of isolated NTD, ID, TRD and CTD domains confirmed their lack of stable secondary structure^4^. It has been experimentally demonstrated that the NTD, CTD and TRD domains can undergo a coil to helix transition, with the TRD showing the greatest tendency for helix formation^4^.

To date, two computational studies of MeCP2 have been reported, and both focus on the ordered MBD domain only. Kucukkal and Alexov reported comparative MD simulations of the R133C mutant and wild-type MBD^5^, and Yang et al. studied the effects of Rett syndrome-causing mutations on the binding affinity of MBD to CpG dinucleotides^6^. The scarcity of computational studies is due to the lack of a three-dimensional structure of the full-length protein. Nevertheless, computer simulations have been able to predict the structures of IDPs. For example, using a coarse-grained model, *ab initio* simulations of pKID successfully modeled its coupled folding and binding to KIX^7^, and a combination of homology and *ab initio* modelling provided valuable insight into the three-dimensional structures of intrinsically disordered e7 proteins^8^. In this work, we used the known structure for the MBD domain as a starting point with the rest of the protein built by *ab initio* modeling. Using a combination of all-atom and coarse-grained simulations, the folding of the full-length MeCP2 and the conformational ensemble it could sample were studied.

## METHODS

### Ab initio modelling

Modeller^9^ version 9.19 was used to build a model for the full-length MeCP2 protein. Using the BLAST algorithm^10^, we searched the UniProt database^11^ for homologues of MeCP2 with a 3D structure. Unfortunately, the only known structures belong to homologues of the MBD domain, which accounts for only ~17% of MeCP2 amino acids and whose structure has already been determined. Thus, we used the Protein Data Bank 1QK9,^1^ which contains the MBD domain structure, as a template. Twenty different models were generated with Modeller^9^. There was little variation between the different models and thus the first model was chosen as the starting structure for the simulations (Fig. S1). With the aim of having a different starting structure for our coarse-grained simulations, a second model was built by refining the loops of the first model using the loopmodel class in Modeller. There is no structural information on this protein besides its known disorder and the structure of the MBD domain, and thus no quality assessment predictors were used to evaluate the generated models. The evaluation will come from the data obtained during the simulations.

The AlphaFold^12^ model for human MeCP2 (UniProt code: P51608) was downloaded from the database hosted by the European Bioinformatics Institute (https://alphafold.ebi.ac.uk). Three simulations were performed with this model as the initial structure, using the procedure described in the next section.

### All-atom simulations

The following procedure was used in all of the all-atom MD simulations: The initial structure was placed in a dodecahedral box in which the distance from the edges of the box to every atom in the protein was at least 1 nm. The box was solvated with water and 150 mM of NaCl was added to reproduce physiological conditions. Counterions were added to maintain the overall charge neutrality of the system. Simulations were performed using GROMACS 2016.3^13^ with the TIP3P water model^14^ and the Amber99SB*-ILDNP force field^15^. The only exception is the set of five replicas for the ID and TRD domains that were run with the CHARMM36IDPSFF force field that is parameterized specifically for intrinsically disordered proteins^16^. This IDPs-specific force field has been shown to produce good results when compared to other force fields in a recent study of amyloid-β^17^, an extensively studied IDP. Table S1 contains the details of the all-atom simulations: three simulations of Modeller models, three simulations of the AlphaFold model and 12 simulations of sections of the ID and TRD domains of different lengths.

Each system was first energy minimized using the method of steepest descents and pre-equilibrated in the canonical ensemble, i.e., at constant particle number, temperature and volume, for 100 ps. Pre-equilibration was followed by a production run with a time step of 2 fs. The Lennard-Jones potential was truncated using a shift function between 1.0 and 1.2 nm. Electrostatic interactions were calculated using the particle-mesh Ewald method (PME)^18,19^ with a real space cut-off of 1.2 nm. The temperature was set to 310 K with the V-rescale algorithm^20^ and pressure was kept at 1 atm using the Parrinello-Rahman barostat^21^. Bonds involving hydrogens were constrained using the Parallel Linear Constraint Solver (P-LINCS) algorithm^22^.

Some systems (marked as “resized” in Table S1) were moved into a smaller simulation box after an initial run in which the protein became more compact. The final configuration of the initial simulation was placed into a new simulation box in which the distance from the edges of the box to every atom in the protein was again at least 1 nm. The new box was solvated with water, 150mM of NaCl and counterions. Each new system was energy minimized and pre-equilibrated in the canonical ensemble before moving to the production run. All parameters mentioned above were kept the same. Trajectory analysis was performed using Gromacs built-in tools^13^ and MDAnalysis^23,24^.

### Coarse-grained simulations

The intermediate-resolution implicit solvent coarse-grained protein model PLUM^25^ by Bereau and Deserno was used to further explore the conformational landscape of MeCP2. This model represents the backbone with near-atomistic resolution, with beads for the amide group N, central carbon Cα and carbonyl group C’. The side chains are represented by single beads located at the first carbon Cβ of the all-atom model. The N and C’ beads can hydrogen bond through a directional potential which depends on the implicit positions of hydrogen and oxygen atoms within them. The PLUM model has been successfully used to study a variety of scenarios such as the aggregation of polyglutamine^26^, β-barrel formation at the interface between virus capsid proteins^27^, folding of transmembrane peptides^28^, and it has been shown to be able to reproduce the secondary structure of small IDPs involved in biomineralization^29^.

Simulations using this model were carried out in GROMACS 4.5.5^13^ specifically modified to support the PLUM model. All interaction parameters were taken from the original work of Bereau and Deserno^25^. The simulations were run in the canonical ensemble (NVT) with a Langevin thermostat with friction constant *Γ* = *τ*^−1^ and an integration timestep of *δt* = 0.01*τ*, where τ is the natural time unit in the simulation. The reduction in degrees of freedom removes friction and speeds up the motion through phase space and thus this time unit is not equivalent to the time step in an all-atom simulation^25^. Table S2 contains the simulation details.

## RESULTS

### The all-atom protein is largely unstructured

A full-length MeCP2 all-atom protein model, henceforth referred to as MeCP2_1, was simulated for 1,550 ns. The protein started with an extended conformation (Fig. 2) in order to minimize bias towards any particular fold. After an initial simulation of 150 ns, the protein had become more compact and it was moved to a smaller box to increase efficiency. Figure 2 shows a drastic decrease in the radius of gyration (R_g_) during the first 20 ns of the simulation, when it went from 8.35 nm to 4.56 nm. Although R_g_ continued to fluctuate, it never surpassed 5 nm. Moving the protein to a smaller box allowed a reduction in the number of water molecules from ~793,000 to ~120,000 (Table S1).

**Figure 2.**
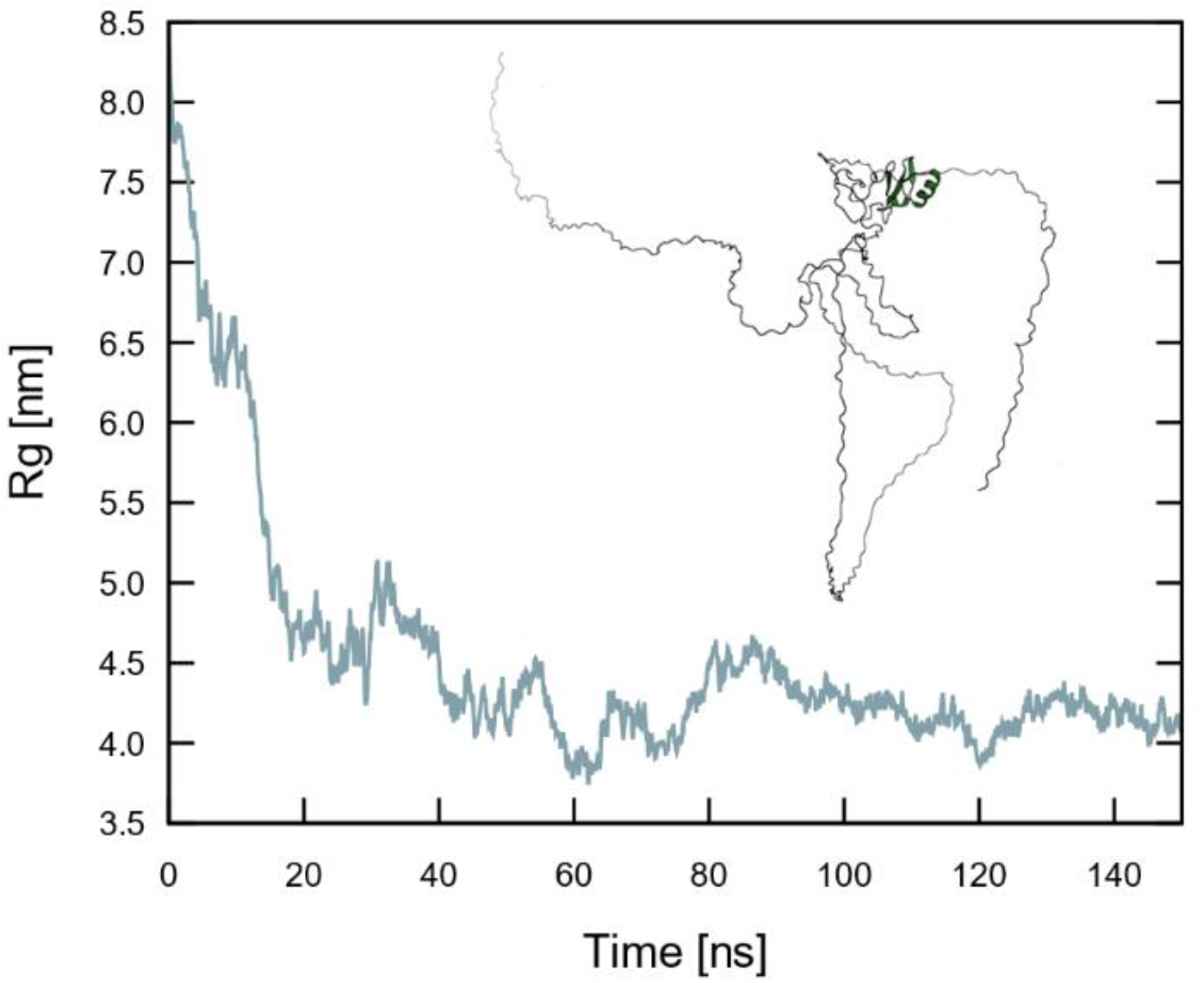
Radius of gyration (R_g_) of the full-length MeCP2 in the initial simulation box. The protein becomes more compact during the first 20 ns. The inset shows the initial structure. MBD, the only ordered domain, is clearly visible.

The protein was then run in the new box for an additional 1,400 ns. The protein remained highly flexible; its root-mean-square deviation (RMSD) from the initial structure continued to show small fluctuations throughout the trajectory, as expected for an IDP (Fig. 3A). The most populated cluster in the last 400 ns of the simulation is largely unstructured with only small motifs of secondary structure (Fig. 3B). Shown in Fig. 3B red are the residues with a root mean square fluctuation (RMSF) larger than 0.6 nm. Figure 3C shows the RMSF of each amino acid throughout the last 400 ns of the trajectory. The residues with the highest RMSF are located in the NTD and CTD-β domains, at the opposite ends of the protein, and in two solvent-exposed loops.

**Figure 3.**
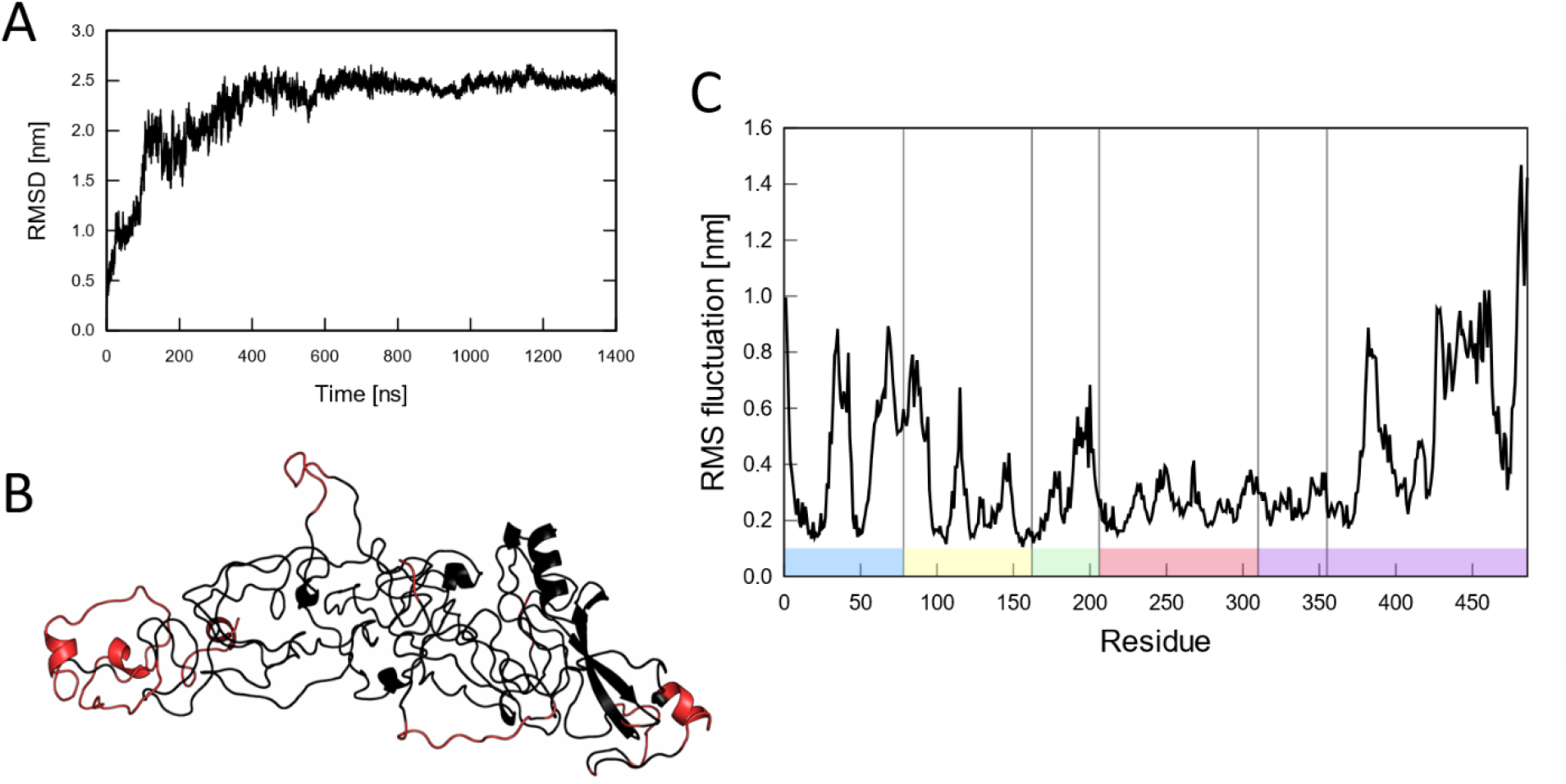
All-atom MD simulation of the full-length MeCP2. A) RMSD of the protein in the smaller simulation box. B) Most sampled cluster throughout the last 400 ns of the simulation. Red: residues with an RMSF higher than 0.6 nm. C) RMSF of the protein throughout the last 400 ns of the simulation. The different domains are marked following the color code in Fig 1.

The last 400 ns of the 1,400 ns trajectory were clustered using the method of Daura *et al*.^30^ with a 0.5 nm cut-off. The secondary structure was computed for the most representative structure in each of the 11 clusters obtained (Table S3). The weighted averages show that 20 residues had an α-helix conformation (4.1%), 113 residues were in β-strands or turns (23.2%) and 338 residues were in random coil (69.7%). This is very similar to experimental data of Adams et al., in particular the amount of α-helix compared to the experimentally (by CD) determined 4%^31^.

We also computed the secondary structure of the protein throughout the last 400 ns of the simulation using DSSP^32,33^. The secondary structure elements in the MBD domain are very stable, appearing in at least 80% of all frames (Fig. 4). Adams *et al*^31^ reported the secondary structure for the MBD domain on its own to be 10% α-helix, 51% β-strands or turns and 38% unstructured, and the NMR structure (PDBid: 1QK9^1^) contains 12% α-helix, 20% β-strands or turns and is 69% unstructured. The MBD domain in our simulation had 15% α-helix, 28% β-strands or turns, and is 57% unstructured. Overall, our simulation is in good agreement with the experimental data. The most disordered domains are the ID domain (Fig. S4) and the CTDα domain (Fig. S6). The NTD domain has two short β-strands and two helices, with one of them present in 60% of the simulation frames (Fig. S3). The TRD domain formed a helix in residues 241 to 244 in 70% of the simulation and two β-strands were observed in 6% of the frames (Fig. S5). The residues in the α-helix correspond to 4% of the TRD residues and the unstructured residues to 87% of the TRD amino acids. This is in good agreement with the 3% of α-helix and 85% of unstructured residues measured by Adams *et al*^31^. Five helices are observed in the CTDβ domain, with two of them present 80% of the simulation (Fig. S7).

**Figure 4.**
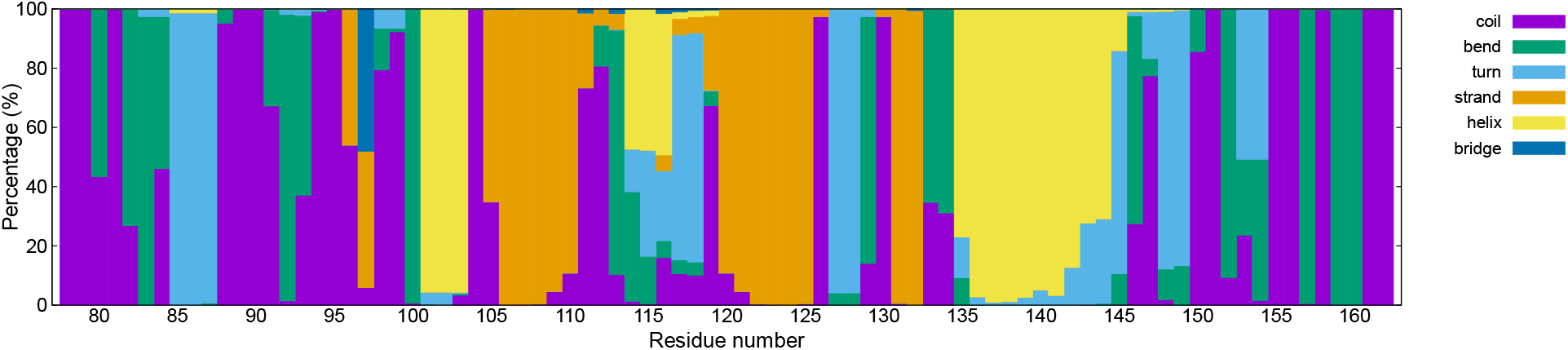
Percentage of frames in the last 400 ns of the MBD domain with every type of secondary structure. The secondary structure of the MBD domain observed in the MeCP2_1 simulation, 15% α-helix, 28% β-strands or turns, and 57% unstructured, is in good agreement with experimental data^31^.

Even though some of the secondary structure elements appeared in only a small fraction of the frames, these could become stable upon interaction with another protein, DNA or a small molecule. In fact, most domains in MeCP2 can bind to DNA; the MBD domain binds to symmetrically methylated 5′CpG3′ pairs with a preference for A/T-rich motifs^34,35^, an autonomous DNA binding domain has been identified in the ID domain^36^, the TRD domain possesses a non-specific DNA binding site^31,36^ and there is a distinct non-specific binding site for unmethylated DNA in the CTDα domain^36^.

Principal component analysis (PCA) of the trajectory underlines the structural rearrangements that the protein undergoes during the first 600 ns of the simulation (Fig. S8A). After this time, the protein explores a much smaller portion of the conformational space. In contrast, the TRD domain begins to sample more conformational space in the second half of the simulation (Fig. S8B).

The experiments by Ghosh *et al*.^37^ showed that the single tryptophan of MeCP2, which is located at position 104 in the MBD domain, is strongly protected from the aqueous environment. Using the STRIDE web server^38^, we computed the relative solvent accessible surface area (rSA) of residue W104 in the four most populated clusters (Table S4). The first four clusters contain 98% of all frames in the last 400 ns simulation. Although there is no consensus on where to set the threshold to determine if an amino acid is buried, it is typically set between 10% and 20%^39,40^. The weighted average for the four clusters gave a rSA of 8.1% and thus it can be considered to be buried inside the protein, in agreement with the experimental data.^37^

R133C is one of the most common disease-causing mutations in the MBD domain^37^. The x-ray structure of an MBD-DNA complex has revealed that Arg 133 is involved in the DNA interaction surface^41^, and the study by Lei *et al*.^42^ found that this residue, together with Arg 111, forms hydrogen bonds with DNA. In order to see if these two residues are solvent accessible in our simulation, we computed their rSA (Tables S5 and S6). Residue R111 had a rSA of 12.7% in the most populated cluster, which can be considered to be buried. However, this amino acid had a high rSA value in the second most populated cluster, giving a weighted average of 20.4%. Therefore, this residue is actually solvent accessible. Residue R133 had a weighted average rSA of 53.7% and thus is also solvent accessible. Kucukkal and Alexov^5^ reported an average number of hydrogen bonds with water of 1.68 for residue R133 and 0.47 for residue R111 in their MBD-only simulations. We obtained an average of 2.96 for R133 and 1.59 for R111 in the last 800 ns of the simulation. It is thus evident, that these residues are more solvent accessible when the full-length protein is considered. Kucukkal and Alexov^5^ did not report the total number of salt bridges observed in their simulations, however, they reported the loss of two salt bridges (R133-E137 and K119-D121) upon mutation of residue R133. We computed all salt bridges in the same manner as them, using the Salt Bridges plugin for VMD^43^. A total of 499 salt bridges were identified but most of them appeared in only a small fraction of the frames and only 35 were stable during the last 400 ns of the simulation (Table S7). Most of these salt bridges occur between the NTD and the MBD domains. Salt bridge K119-D121 is only present in very few frames (Fig. S9A). Lys119 formed hydrogen bonds with neighbouring residues 115-117 and Asp121 with Lys109 and Arg111. The salt bridge R133-E137 can be observed at the beginning of the simulation but is lost in the last 400 ns of the simulation. This is consistent with the study by Kucukkal and Alexov^5^ who observed this salt bridge in their 220 ns simulation (Fig. S9B), but it underlines the need for sufficiently long sampling times.

### Coarse-grained simulations sample two different conformations

In order to investigate other possible folds of the protein, we ran four coarse-grained simulations using the PLUM model^25^. Three different configurations were used as starting points: A) the structure of the all-atom simulation after 800 ns, B) the initial structure built with Modeller, and C) model “B” with its loops refined (Fig. 5).

**Figure 5.**
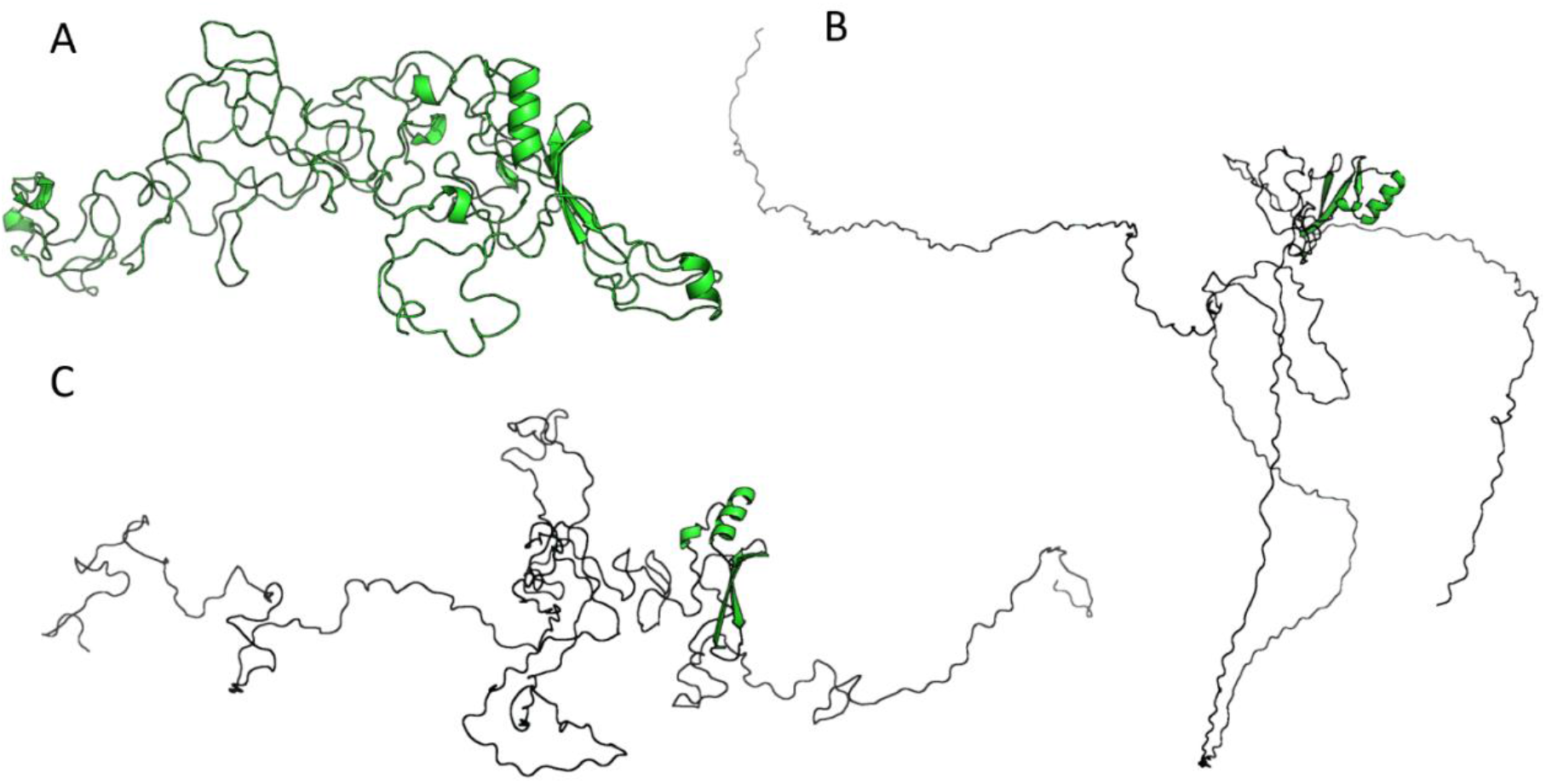
Coarse-grained simulations of MeCP2 using the PLUM model^25^. Simulations started from three different conformations: The end structure of the all-atom simulation after 1150 ns (A), the initial structure built with Modeller^9^ (B) and structure “B” with refined loops (C).

**Figure 6.**
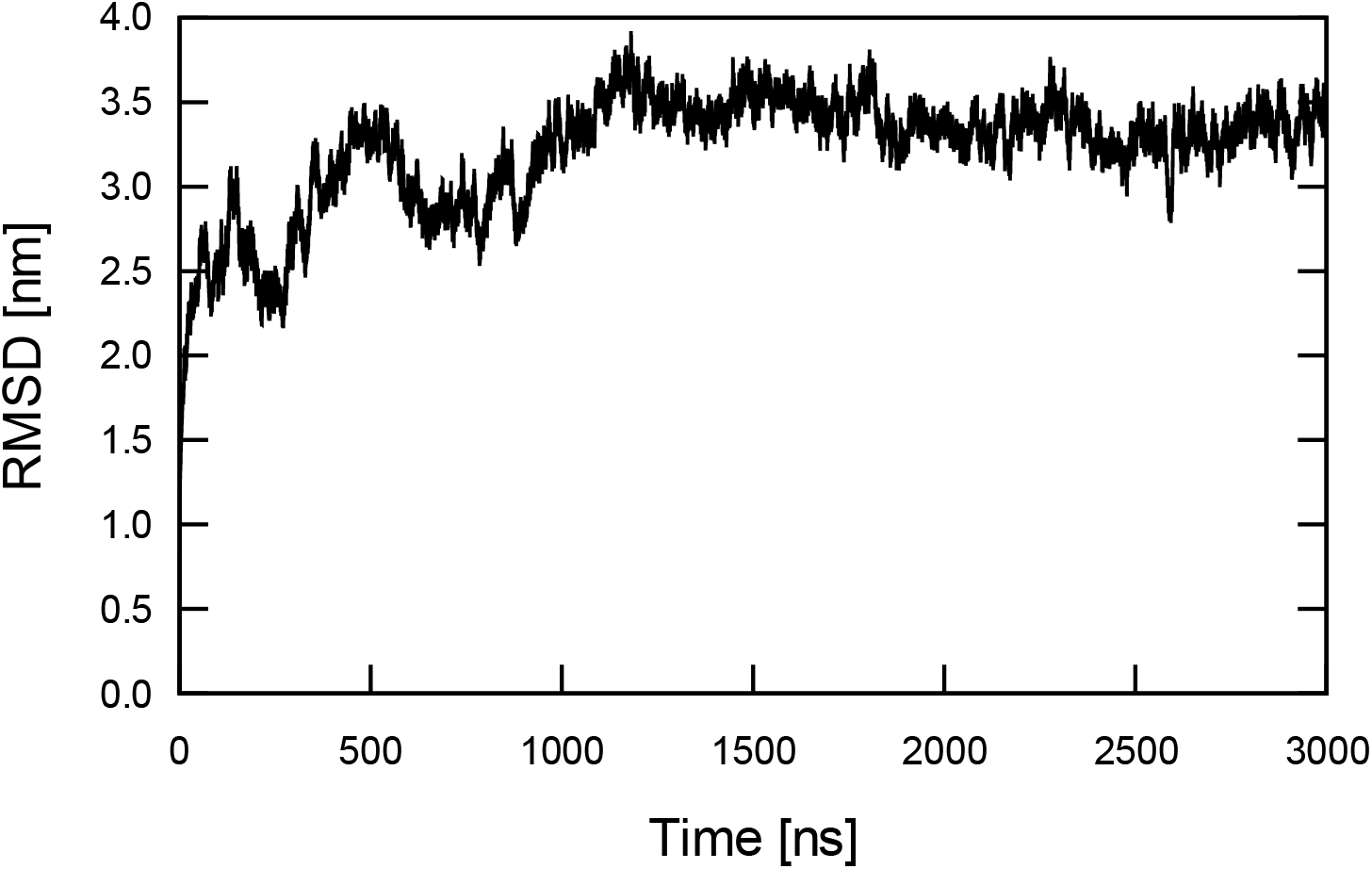
RMSD from the initial structure of the protein (the MeCP2_1 model after 800 ns of simulation) in the CG1 PLUM simulation.

The configuration at 800 ns in the MeCP2_1 simulation (Fig. 5A) was used as the starting point for a coarse-grained simulation, henceforth referred to as CG1. Similar to the RMSD in the all-atom simulation, the RMSD of the protein converges to 3.5 nm but continues to fluctuate. The large RMSD value indicates that the overall topology of the structure changed. A cluster analysis was used help to identify the differences.

We clustered the conformations sampled in the entire trajectory using the method of Daura *et al*.^30^ with a 2.0 nm cut-off. Two main conformations were revealed: 1) a single globule and 2) two globules connected by a loop (Fig. 7). The first two clusters had a single globule configuration but the third had two distinct globules connected by a loop. In this structure, the connecting loop starts at residue 228 and ends in residue 242. A similar conformation can be observed in the eight cluster.

**Figure 7.**
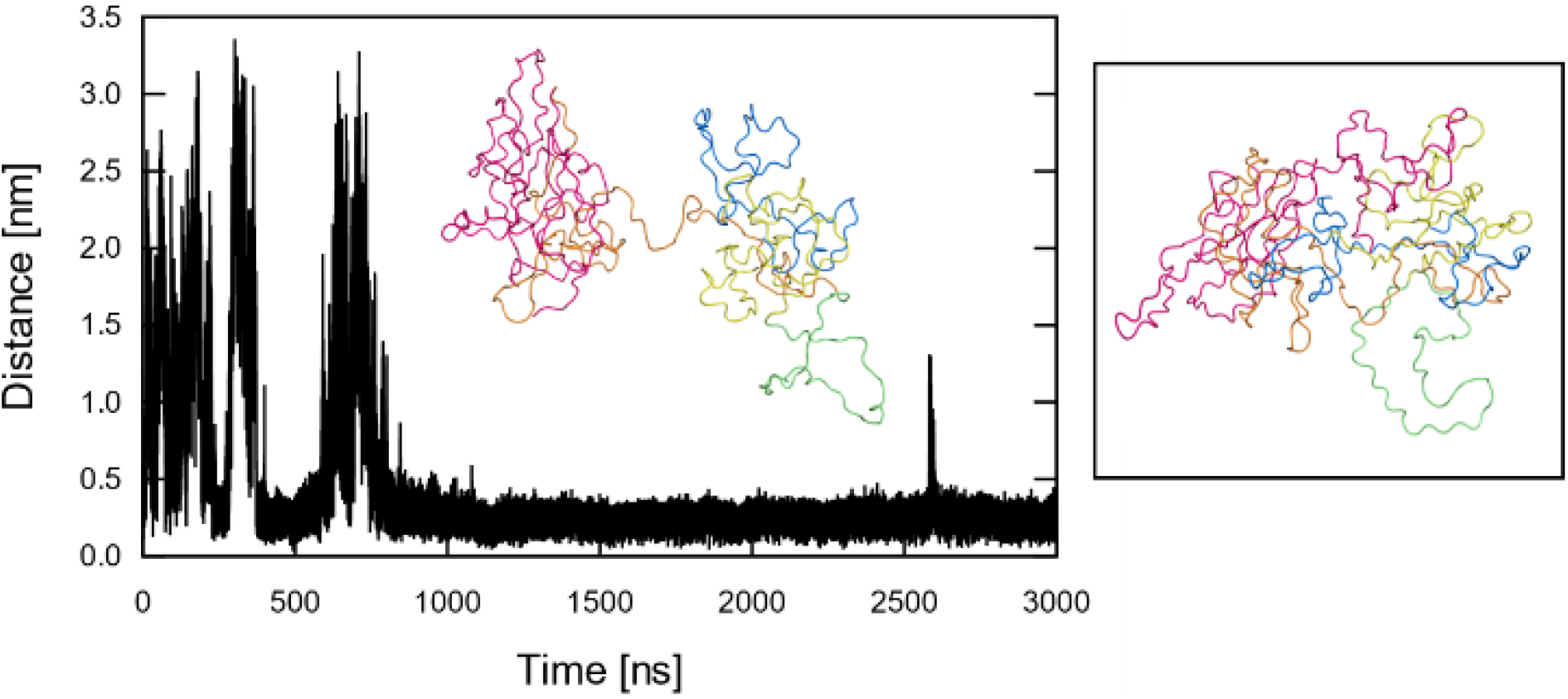
Minimum distance between the two globules in the CG1 PLUM simulation. A two-globule conformation is sampled at the beginning of the simulation at 700 ns and once again at 2,600 ns. The single globule and two-globule conformations have their different domains marked following the color code in Fig. 1.

The minimum distance between the amino acids of the two globules throughout the simulation shows that the two-globule conformation was sampled at the beginning of the simulation, around 700 ns and at 2,600 ns of simulation. The first two times this conformation was sampled, the linker between the two globules was long enough to stabilize it for ~100 ns. In contrast, the two-globule conformation sampled at 2,600 ns had a shorter linker and it coalesced into a single globule after 20 ns.

Two different replicas (simulations CG2 and CG3) were run for the model built with Modeller (Fig. 5B). Their RMSD converged in the first 200 ns but the simulations were extended to 500 ns (Fig. 8A). Since the reference structure for the RMSD calculation is the initial frame i.e., the unfolded structure, a high RMSD value is to be expected. Simulation CG2 sampled conformations similar to those observed in the all-atom MeCP2_1 simulation, albeit more compact (Fig. 8B). Simulation CG3 collapsed into a globule which appears to be an energetic minimum since the system could not sample any other conformations (Fig. 8A). Interestingly, the loop that remained solvent-exposed for this entire simulation, spans residues 168 to 201 and corresponds to the ID domain (Fig. 1).

**Figure 8.**
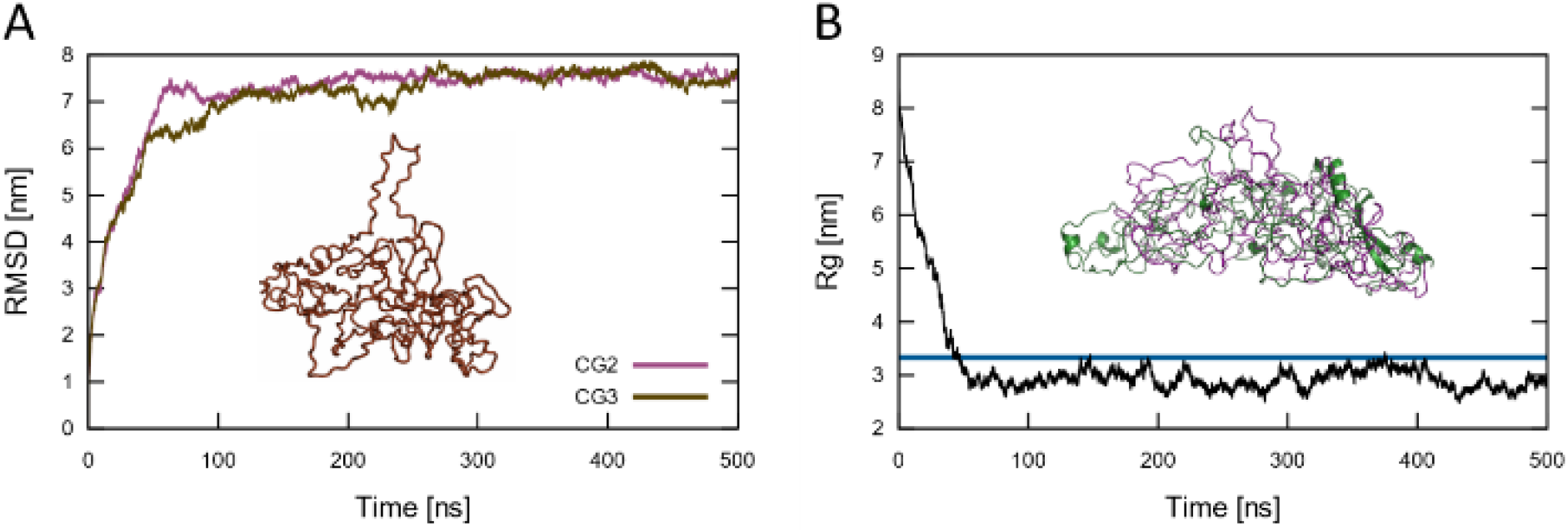
(A) RMSD from the initial structure (Modeller model) of the CG2 and CG3 PLUM simulations Brown: Most populated cluster in the entire CG3 PLUM simulation. (B) Radius of gyration of the CG2 PLUM simulation. Blue line: The average Rg of the MeCP2_1 system. Green: The average structure of the most populated cluster in the MeCP2_1 simulation. Magenta: The average structure of the second most populated cluster in the last 400 ns of the CG2 PLUM simulation.

A fourth coarse-grained simulation (CG4) started from the Modeller model with its loops refined (Fig. 5C). Since the starting structure had not been energy minimized, a high RMSD value is to be expected. This simulation sampled two-globule conformations similar to those observed in the CG1 simulation albeit with the connecting loop located between residues 161 and 205. Interestingly, the location of this loop matches the ID domain (Fig. 1). The protein underwent two main transitions during the simulation. It became more compact during the first 40 ns, it sampled two-globule conformations from 40 to 315 ns, and it sampled a single globule for the rest of the simulation (Fig. 9).

**Figure 9.**
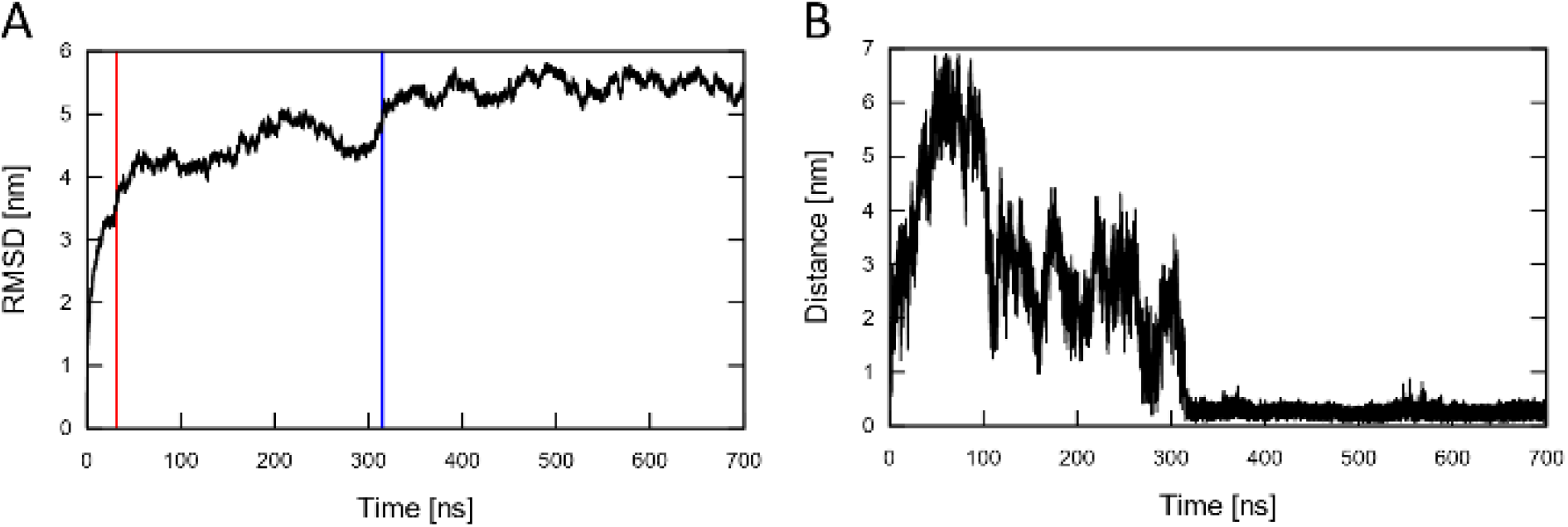
(A) RMSD from the initial structure (Modeller model with refined loops) and (B) minimum distance between the two globules of simulation CG4. The protein becomes more compact and at 40 ns (red line) it starts to sample two-globule conformations. After 270 ns (blue line) the two globules merge together and a single globule is sampled.

A cluster analysis with the method of Daura *et al*.^30^ and a 2.5 nm cutoff of all coarse-grained trajectories concatenated found 23 different clusters (Table S8). The first eight clusters contain 95.3% of all structures sampled. Four of these clusters are single globules and four are two-globule conformations. Only the single globule conformations had overlap between trajectories.

To further understand the conformational space sampled by all coarse-grained trajectories, we performed a single Principal Components Analysis (PCA) on all simulations. Even though each simulation sampled different conformations, these get closer to one another over time, when projected onto the first two eigenvectors (Fig. 10). This implies that the protein tends toward a similar, limited conformational ensemble in all four simulations.

**Figure 10.**
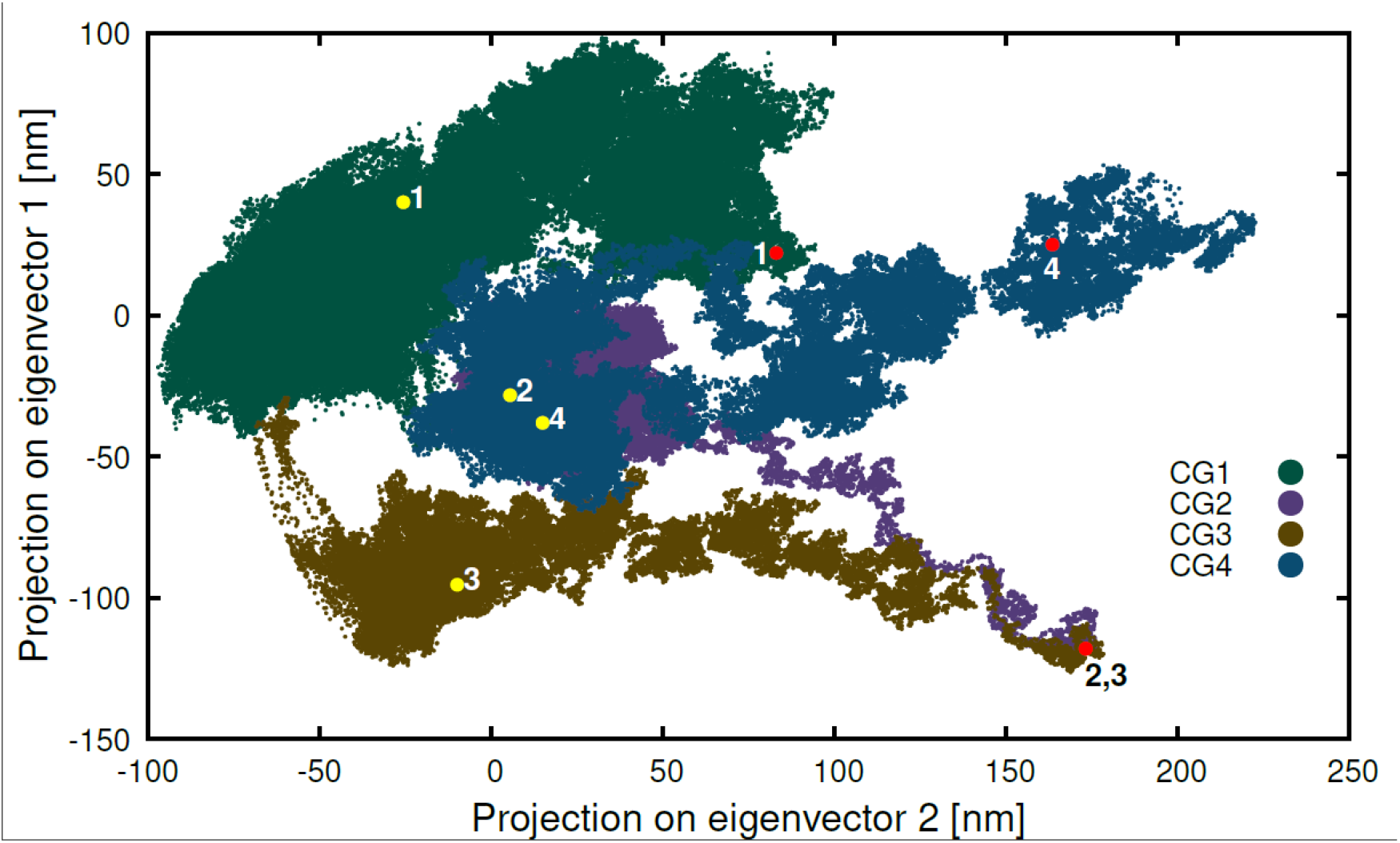
Principal Component Analysis. Projection of all coarse-grained trajectories on the first two eigenvectors, each trajectory is depicted with a different colour. Simulations CG2 and CG3 started from the same conformation. The starting points of all simulations are marked in red and the end points in yellow.

### All-atom two-globule conformations transition into a single globule

Given that the two-globule conformation had only been sampled by the coarse-grained force field, new all-atom simulations were run starting from this conformation. Modeller^9^ was used to generate the initial structures via homology modeling. Three templates were used to generate the models: One for the first globule (NTD and MBD domains), one for residues in the connecting loop (ID and TRD domains) and one for the second globule (CTD domain).

Model MeCP2_2 was built using the two globules from the most populated cluster with a two-globule conformation in the first 500 ns of simulation CG1. The first template contained residues 1-235, the second template had an extended peptide with residues 230-249, and the third one contained residues 311-486 from the second globule (Table S9). The peptide used in the second template was generated using Pymol^44^. Using a longer peptide for the second template produced single-globule models, with the two globules merged into one and a long loop forming a hoop.

In order to study whether the secondary structure in the MBD domain would have any impact on the stability of the two-globule conformation, we generated another model using an all-atom configuration as a template for the first globule. We used the final structure after 400 ns of simulation as the first template for model MeCP2_3. The loop in Model MeCP2_2 was used as the second template and the second globule (residues 311-486) from simulation CG1 as the third template (Table S9).

Model MeCP2_2 collapsed into a single globule after only 20 ns of simulation and did not sample any other conformations, therefore, we did not continue the production run beyond 60 ns. Model MeCP2_3 did not dwell into the same local minimum and its production run was extended to 600 ns. It sampled the two-globule conformation for a longer time but eventually the two globules melted into one, albeit with a more extended structure than the previous model and retaining the secondary structure (Fig. 11). It is possible that this conformation was not stable enough because the connecting loop was not in a water-soluble conformation. Simulations of the connecting loop could help us shed some light on this matter.

**Figure 11.**
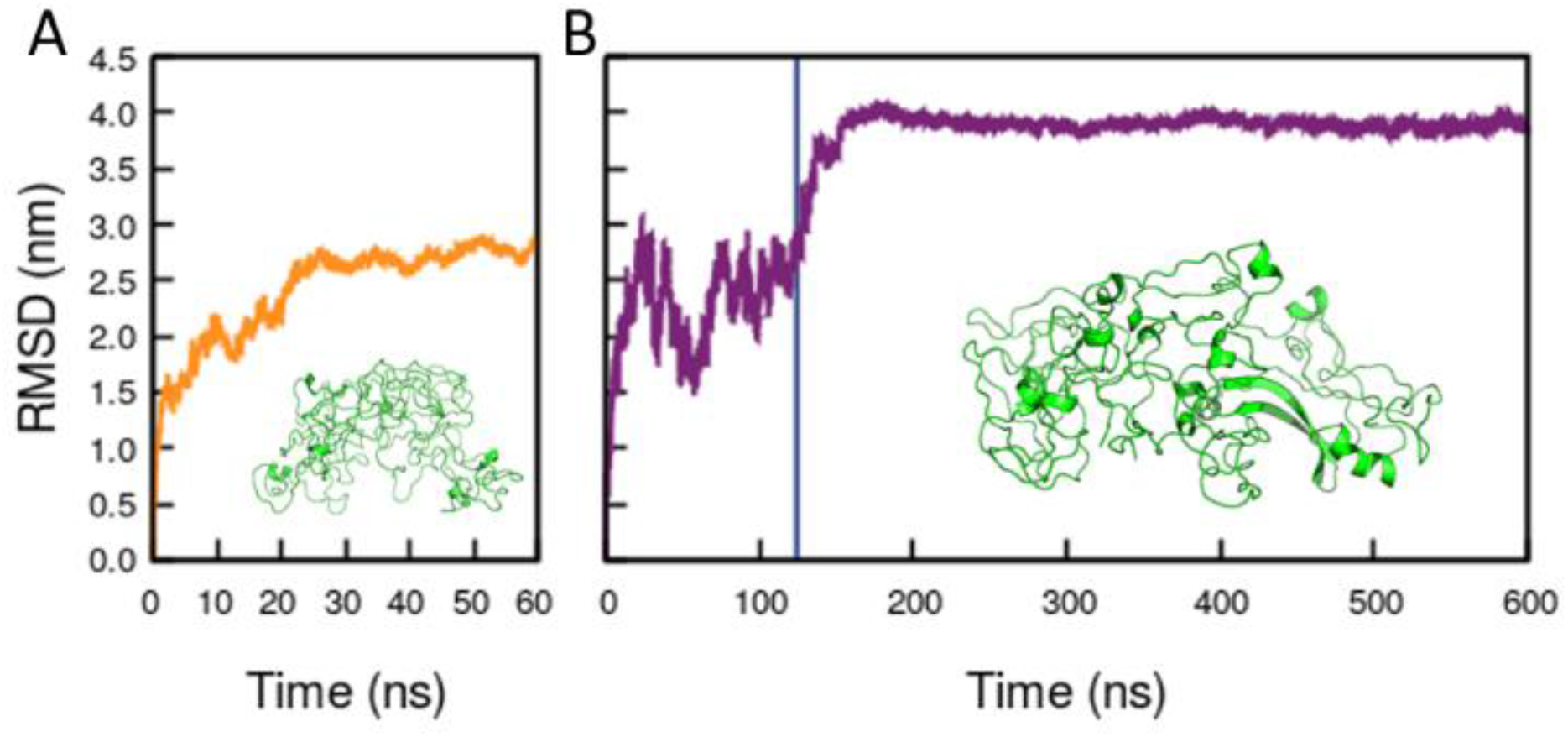
RMSD to the initial structure of models (A) MeCP2_2 and (B) MeCP2_3. Green: The most populated cluster in each trajectory. Model MeCP2_2 samples two-globule conformations during the first 125 ns (blue line), it then undergoes a transition to a single globule.

### Comparing all simulations

Using the PLUMED plugin^45^ for GROMACS^13^, we computed the α-helical content of the all-atom simulations, as well as acylindricity and asphericity of all simulations.

The α-helical content was computed by generating a set of all possible six residue sections in the protein and calculating the RMSD distance between each residue configuration and an idealized α-helical structure. This is done by calculating the following sum of functions of the RMSD distances,

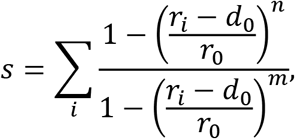

where the sum runs over all possible segments of an α-helix. This collective variable was first defined by Pietrucci and Laio^46^ and all parameters were set equal to those used in their original paper: *d*_0_ = 0.0, *r*_0_ = 0.08 nm, *n* = 8 and *m* = 12.

Model MeCP2_3 sampled conformations with a wider array of values for both the α-helical content and the R_g_ than the MeCP2_1 simulation (Fig. 12). The trajectory analyzed for this all-atom simulation does not include the 150 ns from the bigger simulation box in which the protein underwent initial folding.

**Figure 12.**
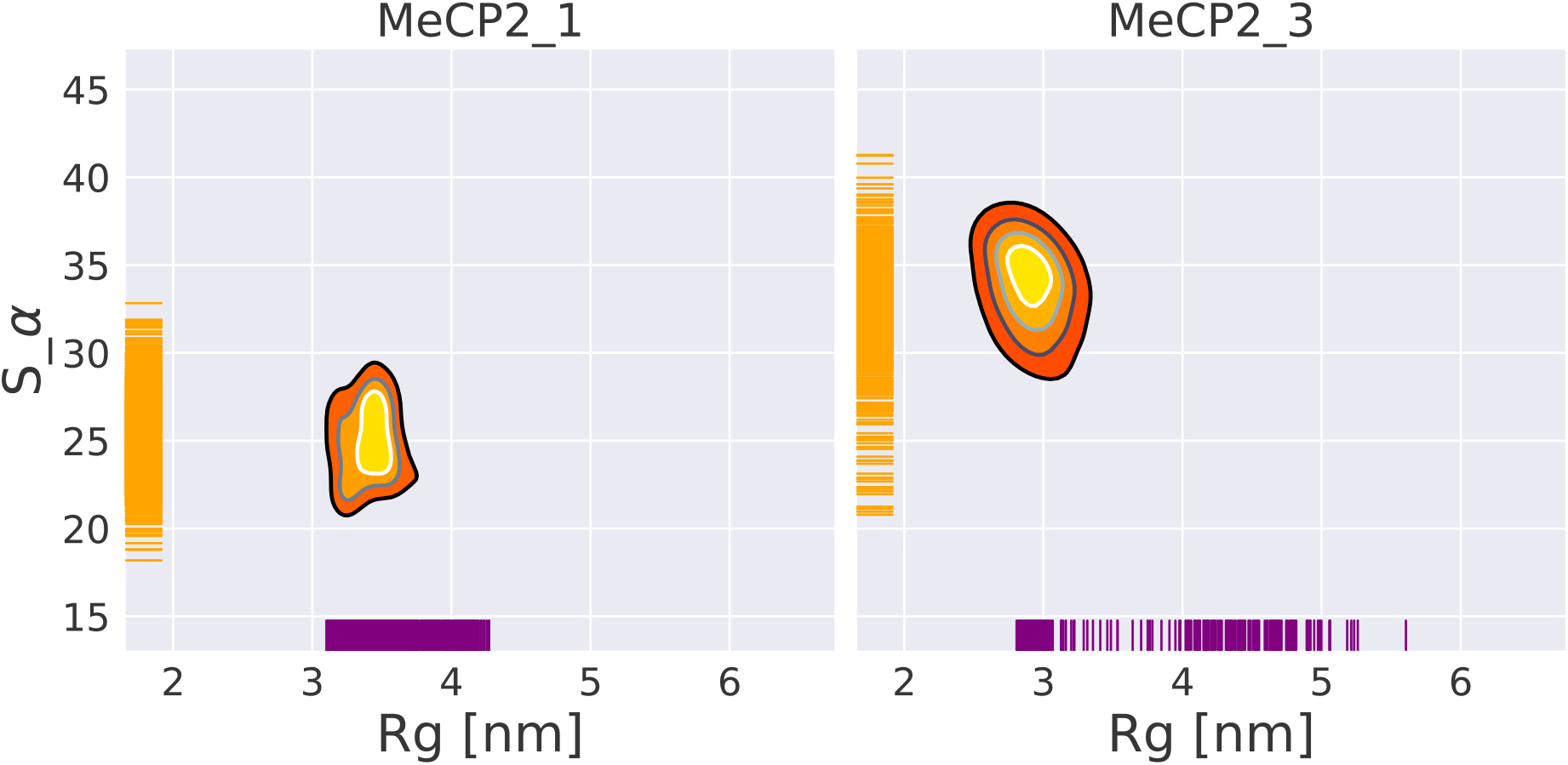
α-helical content of the protein structure vs radius of gyration in the all-atom and two-globule all-atom simulations. Comparison between the all-atom simulation (MeCP2_1, left) that started from an extended structure and the one that started from a two-globule conformation (MeCP2_3, right). Orange: Individual measurements of α-helical content. Purple: Individual measurements of R_g_.

In 1971, Šolc showed that the shape of polymers can be quantified using the eigenvalues (L_1_, L_2_ and L_3_) of the tensor of gyration^47^. The symmetry of a polymer, or in this case, of a peptide, can be described by asphericity,

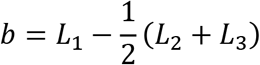

and acylindricity,

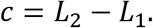

Figure 13 shows the results. All simulations sampled similar values; however, the coarse-grained simulations sampled a wider array of values. From the four coarse-grained simulations, simulation CG1 is the most akin to the all-atom simulations.

**Figure 13.**
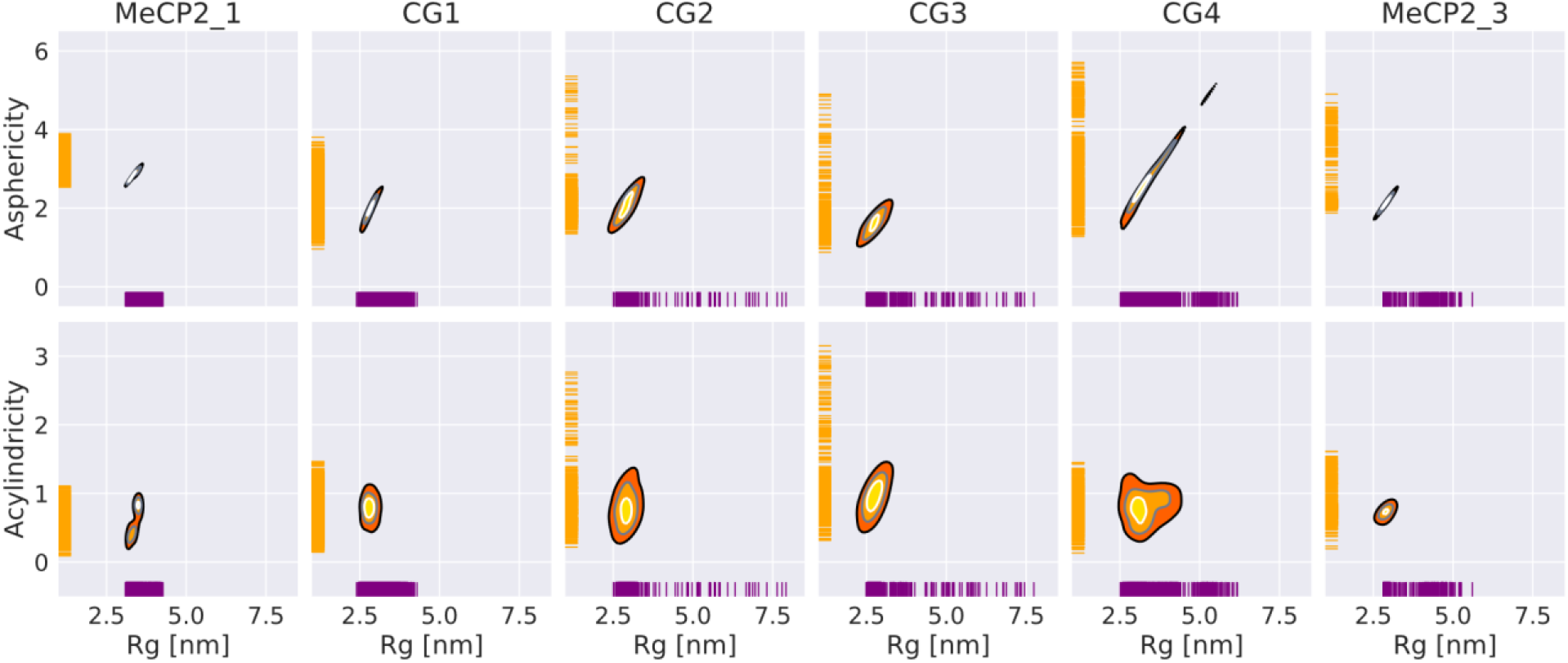
Acylindricity and asphericity vs radius of gyration in the all-atom, coarse-grained and two-globule all-atom simulations. The coarse-grained simulations sampled structures with lower asphericity, higher acylindricity and higher radius of gyration than the all-atom simulations. Orange: Individual measurements of asphericity and acylindricity. Purple: individual measurements of radius of gyration.

### The ID and TRD domains are highly flexible

In order to thoroughly explore the conformations that the flexible ID and TRD domains that form the connective loop can sample, all-atom simulations were run on the ID and TRD domains (residues 164-310). Five replicas were run with two different force fields: Amber99SB*-ILDNP^15^ (simulations A1-A5 in Table S1) and CHARMM36IDPSFF^16^ (simulations C1-C5 in Table S1), using the loop in model MeCP2_3 as the initial structure.

Figure 14 shows R_g_ and end-to-end distance of all structures sampled by the ten simulations. One of these simulations sampled very compact structures but the other nine sampled an array of structures with end-to-end distances between from 3 nm to 23 nm, and radius of gyration from 2.5 nm to 6.5 nm. Table S10 shows the most sampled conformations in all ten simulations. Overall, the Amber force field sampled more compact structures than the Charmm force field, in agreement with previous studies^48–50^.

**Figure 14.**
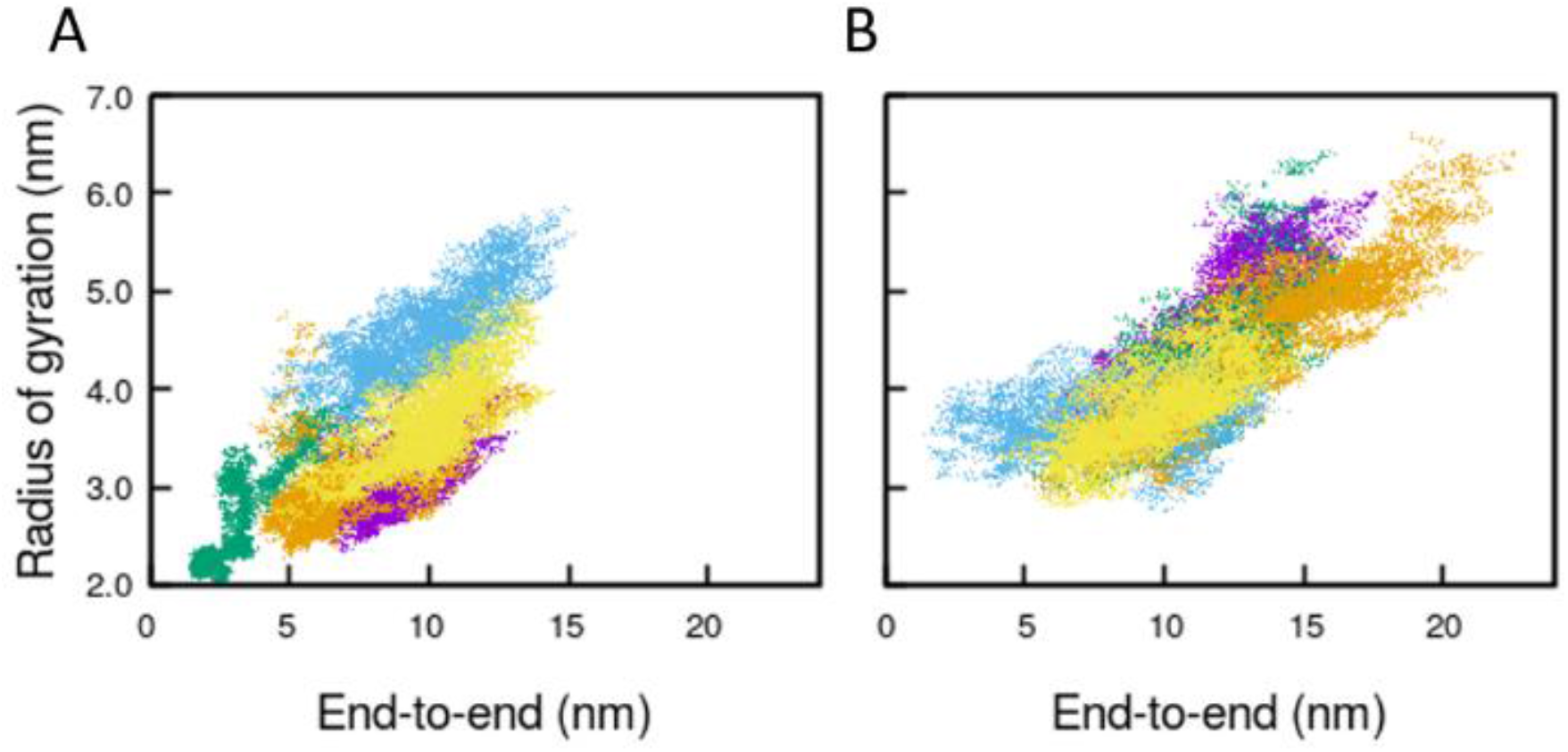
Radius of gyration vs end-to-end distance of five all-atoms simulations of the ID and TRD domains run with Amber99SB*-ILDNP (A) and CHARMM36IDPSFF (B). The peptide is unstructured and can sample a large number of conformations, from compact (low radius of gyration and end-to-end distances) to extended structures (large end-to-end distances). Each simulation is shown in a different color.

In order to understand the role of length in the connecting loop between globules, we simulated the two connecting loops found in the coarse-grained simulations. Two all-atom simulations were performed, one in which the loop spanned residues 228 to 242 (observed in simulation CG1, see Table S2), and another with the loop containing residues 161 to 205 (observed in CG4, see Table S2). The initial structures were taken from the most representative structure of the two-globule conformation in the corresponding coarse-grained trajectory. Modeller^9^ was used to add the missing side-chains and to obtain all-atom structures. The shorter loop (residues 228-242) sampled conformations with the radius of gyration lower than 1.5 nm, whereas the longer loop (residues 161-205) had conformations with the radius of gyration of up to 2.5 nm (Fig. 15).

**Figure 15.**
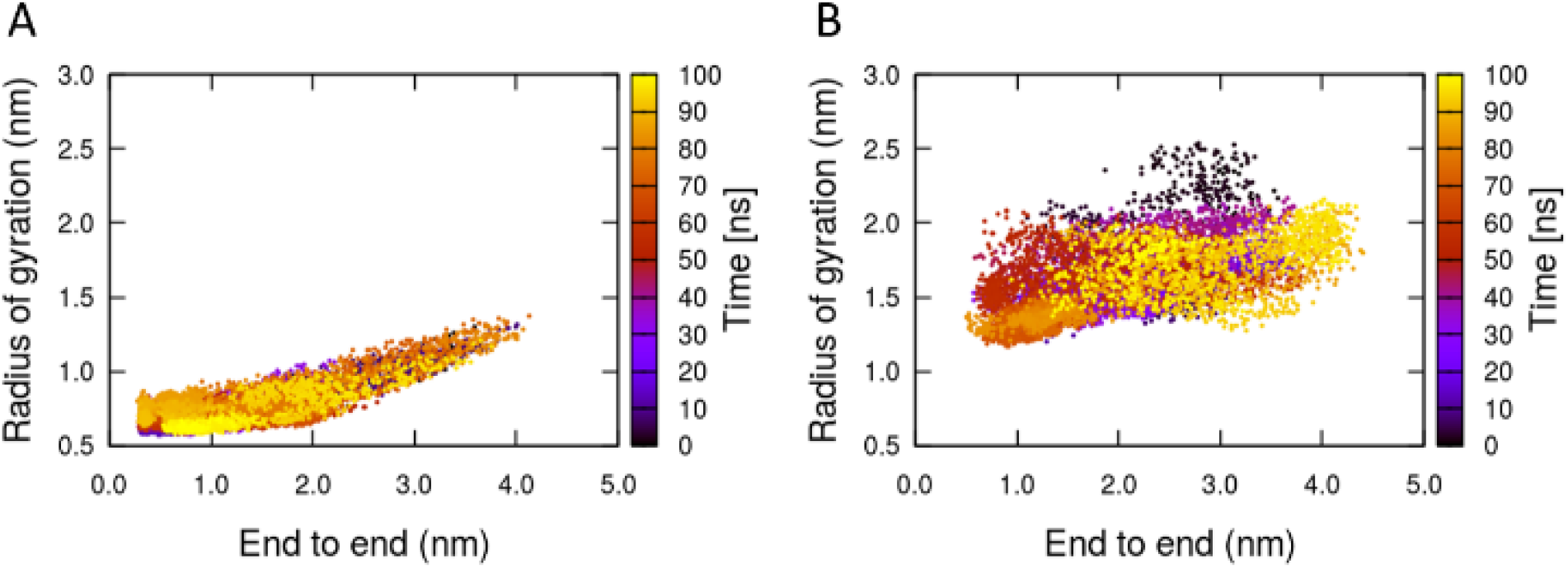
Radius of gyration vs end-to-end distance in all-atom simulations of the two connecting loops found in the coarse-grained simulations. A) Loop with residues 228-242. B) Loop with residues 161-205. The shorter loop sampled more compact structures.

Comparing these two simulations with those of the entire ID and TRD domains (Fig. 14) further underlines the relationship between the length of the loop and its compactness for these particular sequences and range of lengths. The shorter loops observed between the two globules may result in insufficient spacing to stabilize the two-globule conformations sampled in the coarse-grained trajectories, which would explain why they eventually merged into a single globule. Moreover, the two-globule all-atom simulations may be unstable due to the poor initial conditions of the loop generated with Modeller. We hypothesize that a stable two-globule conformation would feature a longer separating loop than observed in our simulations.

### Comparing the simulations with AlphaFold prediction

Last year, the field of bioinformatics had a major breakthrough when the deep learning model AlphaFold was able to successfully predict the three-dimensional structure of proteins from their sequence^12^. Since then, the model has been used to predict 98.5% of the proteins in the human proteome^51^; all the structures are available to the community in a database hosted by the European Bioinformatics Institute (https://alphafold.ebi.ac.uk). Nevertheless, predicting the structure of IDPs remains a challenge, as the vast number of low and very low confidence regions from the structures predicted by AlphaFold overlap with regions predicted to be disordered^52^.

The model we built with Modeller^9^ (MeCP2_1) has the N- and C-terminal ends extended into the solvent, and its radius of gyration is large (8.4 nm). In contrast, the model predicted by AlphaFold^12^ is much more compact (*R_g_* = 4.9 nm) and with an overall spherical shape (Fig. 16). The per-residue confidence score (pLDDT) of almost all residues is either low or very low; only the MBD domain was predicted with confidence (pLDDT > 70). Since the model had such low confidence, we used it as the initial structure for three all-atom MD simulations (Table 1).

**Figure 16.**
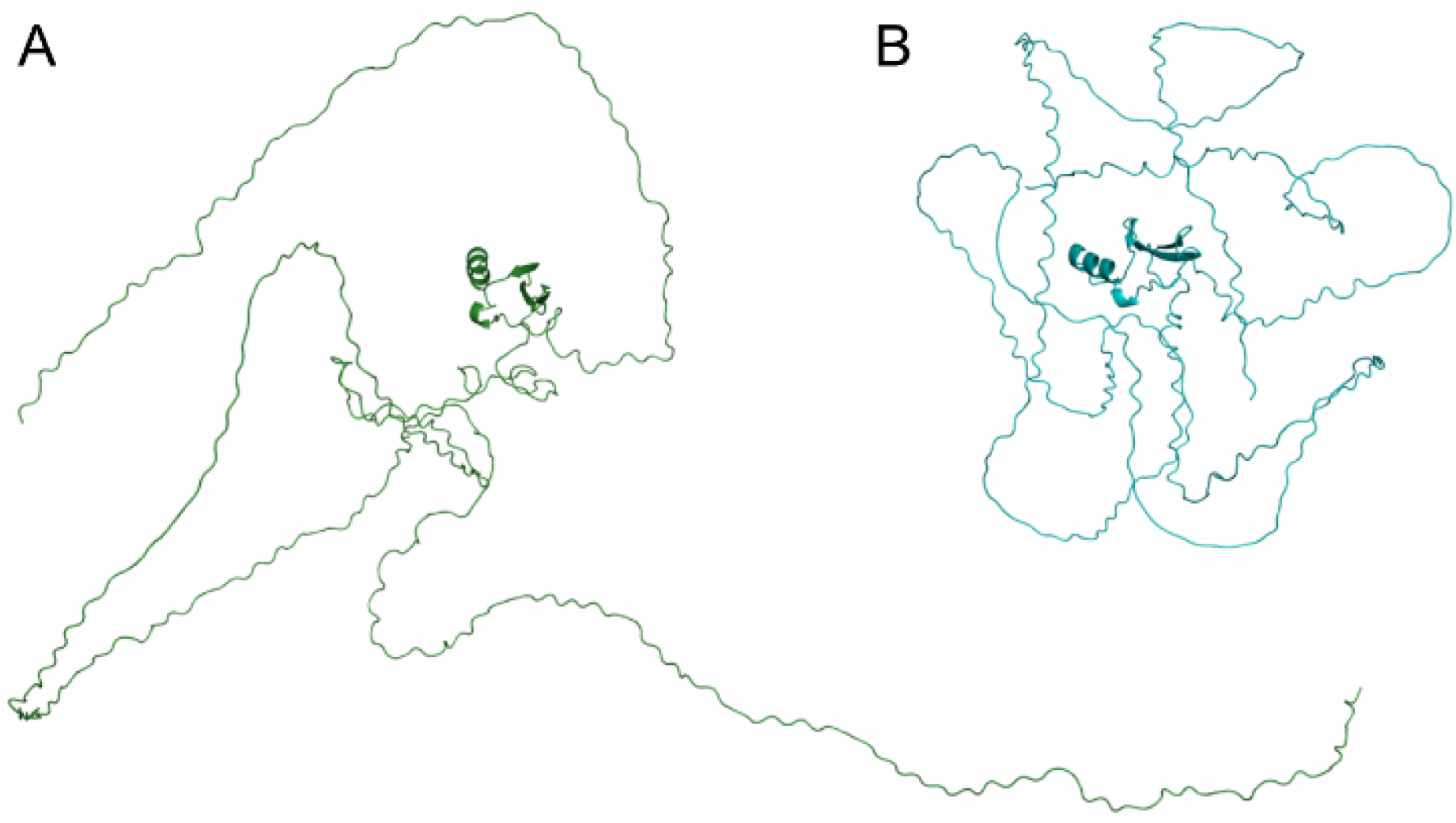
Comparison of the models generated by A) Modeller^9^ and B) AlphaFold^12^, with their respective MBD domains aligned with each other. The model generated by AlphaFold is much more compact than the one generated by Modeller.

Three replicas of the AlphaFold^12^ model were run for 400 ns each. Each simulation sampled a different folding path and converged to a different conformation (Fig. S10). Using the PLUMED plugin^45^ for GROMACS^13^, their α-helical content, acylindricity and asphericity were determined (Fig. S11). The conformations sampled by all three AlphaFold^12^ simulations have similar asphericity and acylindricity values. They sampled conformations that are more spherical and less cylindrical than the conformations sampled by the MeCP2_1 Modeller^9^ simulation. Since the starting structure has a low radius of gyration (*R_g_* = 4.9 nm) we argue that it introduced a bias in the folding path towards more compact structures. Each AlphaFold^12^ replica had a different α-helical content, and only one of them sampled values to similar those observed in the Modeller^9^ simulation.

The AlphaFold prediction was not stable in water and, the only exception being the MBD domain, underwent further folding of all of its domains. Although AlphaFold does a remarkable job predicting the presence of disorder, it cannot solve IDP structures^53^. These simulations should serve as a cautionary tale on the use of predicted models for IDPs; as explained by Strodel in her review^54^, extensive simulations are recommended to equilibrate the protein and sample its conformational space.

## CONCLUSIONS

In this work we have presented a multiscale study of MeCP2, comprising six all-atom and four coarse-grained simulations of the full-length protein, as well as twelve all-atom simulations of the ID and TRD domains. Together, they represent the first computational attempt to study the full-length MeCP2 protein.

The initial model was built starting from the NMR structure of the MBD domain^1^ and building the rest of the protein by *ab initio* modeling. Two main different conformations were sampled in the coarse-grained simulations: a globular structure similar to the one observed in the all-atom force field and a two-globule conformation. This second conformation was not stable in the all-atom force field, probably because the length of the connecting loop was not long enough to be water-soluble. The conformational ensemble sampled by the 1,550 ns all-atom simulation is in good agreement with the available experimental data^1,31^. Our model had 4.1% of α-helix content compared to 4% found experimentally^31^. In addition, our model predicted amino acid W104 to be buried, and amino acids R111 and R133 to be solvent accessible, in accordance with experiments^37,41,42^. Finally, we used the model predicted by AlphaFold^12^ to run three all-atom simulations. The model was not stable in water and underwent further folding. This model is more compact than the one predicted with Modeller^9^, and consequently, it sampled conformations more compact and spherical than those sampled in our Modeller simulations. We recommend caution when using structures of intrinsically disordered proteins predicted by AlphaFold.

With a total of 3 μs of atomistic simulations and 4.7 μs of coarse-grained trajectories of full-length MeCP2 models, extensive conformational space of this protein was sampled. Our longest atomistic simulation (MeCP2_1) converged after 800 ns to a very stable structure. When compared to CG, it is reasonable to assume that the all-atom models are more accurate, so the drift of the CG models towards more compact structures is likely to be an artifact. The results show that no single method (atomistic or CG simulations, or AlphaFold modelling) is sufficient on its own for predicting the conformational ensemble of a large IDP such as MeCP2. Our simulations add structural and dynamical detail to the low-resolution information previously available from experiments and could help study disease-associated mutations in their structural context.

We finish by speculating on how the one- and two-globule conformations that were observed in CG simulations and also transiently in MD simulations, could be investigated experimentally – as discussed above, IDPs pose formidable challenges to both experiments and simulations. One possible way might be high-speed atomic force microscopy (HS-AFM) that has very recently been demonstrated to be able to characterize the structure and dynamics of IDPs (polyglutamine tract binding protein-1 and four of its variants as well as two other IDPs) by Kodera *et al*^55^. In particular, for some of their systems they reported temporarily appearing two-globule conformations and order-disorder transition with an associated change in the (relatively short) linking intrinsically disorder region between the globules. Given that MeCP2 has a long and very flexible disordered region spanning the ID and TRD domains, it is tempting to speculate that fluctuations between the one- and two-globule conformation might be directly detectable or/and inducible in HS-AFM. This seems feasible since force spectroscopy^56^ and MD simulations^57^ have shown that for intrinsically disordered regions forces in the range of a few tens of pN may cause significant stretching and that the free energy barriers are very low. In HS-AFM, the forces are higher up to about 100 pN and there is frictional interaction, albeit very small, with the substrate^55,58^. Thus, the two-globule state that was only marginally stable in current simulations might also be observable in HS-AFM. Such experiments would potentially also allow investigation of the properties of the linker and the globules.

## ASSOCIATED CONTENT

### Supporting Information

Figure S1: Rejected models of the full-length MeCP2 protein, Table S1: Details of all-atom simulations, Table S2: Details of coarse-grained simulations, Figure S2: RMSD of the TRD domain, Table S3: Secondary structure content of the all-atom MeCP2_1 simulation, Figures S3-S7: Percentage of frames in MeCP2_1 with every type of secondary structure, Figure S8: PCA of the MeCP2_1 simulation, Tables S4-S6: rSA of residues W104, R111 and R133, Table S7: Salt bridges in the MeCP2_1 simulation, Figure S9: Salt bridge interactions the MeCP2_1 simulation, Table S8: Clusters sampled in the coarse-grained simulations, Table S9: Templates used to generate models MeCP2_2 and MeCP2_3, Table S10: Conformations sampled by the ID+TRD domains simulations, Figure S10: RMSD of the three AlphaFold simulations, Figure S11: Asphericity, acylindricity and α-helical content of the protein structure, vs radius of gyration in the AlphaFold models.

## AUTHOR INFORMATION

### Author Contributions

The manuscript was written through contributions of all authors. All authors have given approval to the final version of the manuscript.

## ACKNOWLEDGMENT

CCG thanks the Province of Ontario Trillium Scholarship Program and Mitacs for their Globalink Research Award. MK thanks the Natural Sciences and Engineering Research Council of Canada (NSERC) and the Canada Research Chairs Program. JH acknowledges support from the French National Research Agency under grant LABEX DYNAMO (ANR-11-LABX-0011), and from the Laboratoire International Associé CNRS/UIUC. Computing facilities were provided by SHARCNET (www.sharcnet.ca), Compute Canada (www.computecanada.ca).

## ABBREVIATIONS

MeCP2: Methyl CpG binding protein 2
IDP: intrinsically disordered protein
MBD: methyl-CpG binding domain
TRD: transcriptional repression domain
NTD: N-terminal domain
ID: intervening domain
CTD: C-terminal domain
CD: circular dichroism
MD: molecular dynamics
P-LINCS: Parallel Linear Constraint Solver
PME: particle-mesh Ewald method
RMSD: root-mean-square deviation
RMSF: root mean square fluctuation
PCA: principal component analysis

## FOR TABLE OF CONTENTS ONLY

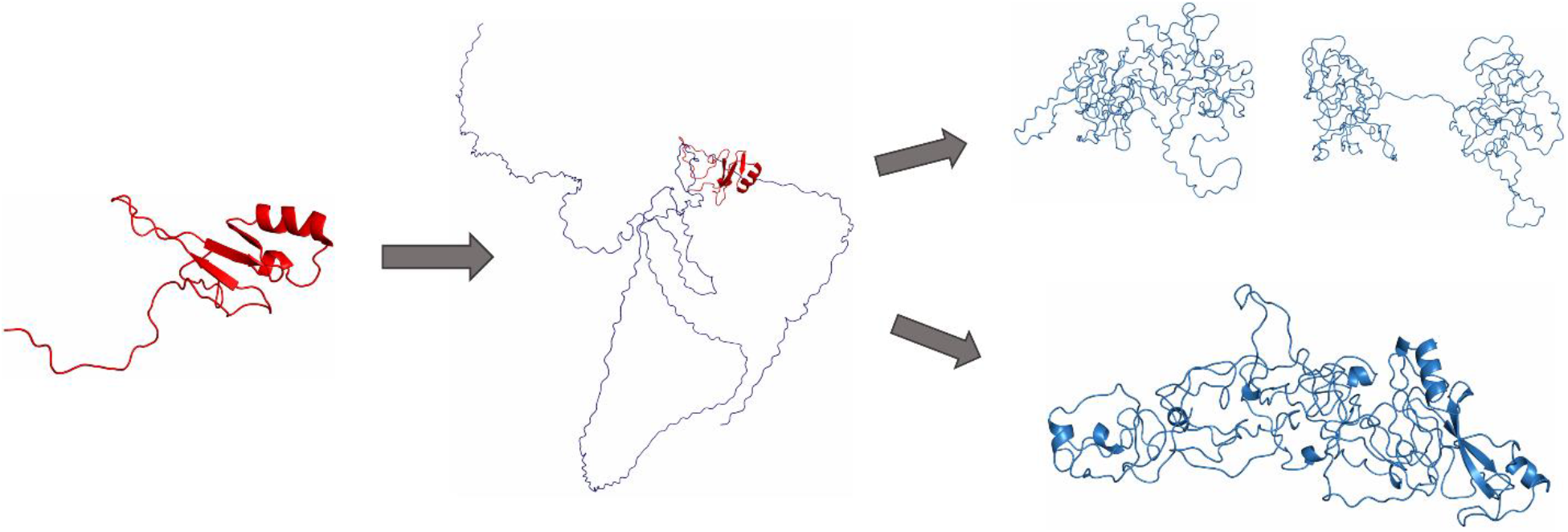

## REFERENCES

(1) Wakefield, R. I. D.; Smith, B. O.; Nan, X.; Free, A.; Soteriou, A.; Uhrin, D.; Bird, A. P.; Barlow, P. N. The Solution Structure of the Domain from MeCP2 That Binds to Methylated DNA. J. Mol. Biol. 1999, 291 (5), 1055–1065. https://doi.org/10.1006/jmbi.1999.3023.

(2) Hite, K. C.; Adams, V. H.; Hansen, J. C. Recent Advances in MeCP2 Structure and Function. Biochem. Cell Biol. 2009, 87 (1), 219–227. https://doi.org/10.1139/O08-115.

(3) Hansen, J. C.; Wexler, B. B.; Rogers, D. J.; Hite, K. C.; Panchenko, T.; Ajith, S.; Black, B. E. DNA Binding Restricts the Intrinsic Conformational Flexibility of Methyl CpG Binding Protein 2 (MeCP2). J. Biol. Chem. 2011, 286 (21), 18938–18948. https://doi.org/10.1074/jbc.M111.234609.

(4) Hite, K. C.; Kalashnikova, A. A.; Hansen, J. C. Coil-to-Helix Transitions in Intrinsically Disordered Methyl CpG Binding Protein 2 and Its Isolated Domains. Protein Sci. 2012, 21 (4), 531–538. https://doi.org/10.1002/pro.2037.

(5) Kucukkal, T. G.; Alexov, E. Structural, Dynamical, and Energetical Consequences of Rett Syndrome Mutation R133C in MeCP2. Comput. Math. Methods Med. 2015, 2015.

(6) Yang, Y.; Kucukkal, T. G.; Li, J.; Alexov, E.; Cao, W. Binding Analysis of Methyl-CpG Binding Domain of MeCP2 and Rett Syndrome Mutations. ACS Chem. Biol. 2016, 11 (10), 2706–2715. https://doi.org/10.1021/acschembio.6b00450.

(7) Kurcinski, M.; Kolinski, A.; Kmiecik, S. Mechanism of Folding and Binding of an Intrinsically Disordered Protein as Revealed by Ab Initio Simulations. J. Chem. Theory Comput. 2014, 10 (6), 2224–2231. https://doi.org/10.1021/ct500287c.

(8) Nicolau-Junior, N.; Giuliatti, S. Modeling and Molecular Dynamics of the Intrinsically Disordered E7 Proteins from High- and Low-Risk Types of Human Papillomavirus. J. Mol. Model. 2013, 19 (9), 4025–4037. https://doi.org/10.1007/s00894-013-1915-8.

(9) Šali, A.; Blundell, T. L. Comparative Protein Modelling by Satisfaction of Spatial Restraints. J. Mol. Biol. 1993, 234, 779–815.

(10) Altschul, S. F.; Gish, W.; Miller, W.; Myers, E. W.; Lipman, D. J. Basic Local Alignment Search Tool. J. Mol. Biol. 1990, 215 (3), 403–410. https://doi.org/10.1016/S0022-2836(05)80360-2.

(11) Leinonen, R.; Garcia Diez, F.; Binns, D.; Fleischmann, W.; Lopez, R.; Apweiler, R. UniProt Archive. Bioinformatics 2004, 20 (17), 3236–3237. https://doi.org/10.1093/bioinformatics/bth191.

(12) Jumper, J.; Evans, R.; Pritzel, A.; Green, T.; Figurnov, M.; Ronneberger, O.; Tunyasuvunakool, K.; Bates, R.; Žídek, A.; Potapenko, A.; Bridgland, A.; Meyer, C.; Kohl, S. A. A.; Ballard, A. J.; Cowie, A.; Romera-Paredes, B.; Nikolov, S.; Jain, R.; Adler, J.; Back, T.; Petersen, S.; Reiman, D.; Clancy, E.; Zielinski, M.; Steinegger, M.; Pacholska, M.; Berghammer, T.; Bodenstein, S.; Silver, D.; Vinyals, O.; Senior, A. W.; Kavukcuoglu, K.; Kohli, P.; Hassabis, D. Highly Accurate Protein Structure Prediction with AlphaFold. Nature 2021, 596, 583–589. https://doi.org/10.1038/s41586-021-03819-2.

(13) James Abraham, M.; Murtola, T.; Schulz, R.; Páll, S.; Smith, J. C.; Hess, B.; Lindahl, E. GROMACS : High Performance Molecular Simulations through Multi-Level Parallelism from Laptops to Supercomputers. SoftwareX 2015, 2, 19–25. https://doi.org/10.1016/j.softx.2015.06.001.

(14) Jorgensen, W. L.; Chandrasekhar, J.; Madura, J. D. Comparison of Simple Potential Functions for Simulating Liquid Water Liquid Water. J. Chem. Phys. 1983, 79, 926–935.

(15) Aliev, A. E.; Kulke, M.; Khaneja, H. S.; Chudasama, V.; Sheppard, T. D.; Lanigan, R. M. Motional Timescale Predictions by Molecular Dynamics Simulations: Case Study Using Proline and Hydroxyproline Sidechain Dynamics. Proteins 2014, 82 (2), 195–215. https://doi.org/10.1002/prot.24350.

(16) Liu, H.; Song, D.; Lu, H.; Luo, R.; Chen, H. F. Intrinsically Disordered Protein-Specific Force Field CHARMM36IDPSFF. Chem. Biol. Drug Des. 2018, 92 (4), 1722–1735. https://doi.org/10.1111/cbdd.13342.

(17) Samantray, S.; Yin, F.; Kav, B.; Strodel, B. Different Force Fields Give Rise to Different Amyloid Aggregation Pathways in Molecular Dynamics Simulations. J. Chem. Inf. Model. 2020, 60 (12), 6462–6475. https://doi.org/10.1021/acs.jcim.0c01063.

(18) Darden, T.; York, D.; Pedersen, L. Particle Mesh Ewald: An N·log(N) Method for Ewald Sums in Large Systems. J. Chem. Phys. 1993, 98 (12), 10089–10092. https://doi.org/10.1063/1.464397.

(19) Essmann, U.; Perera, L.; Berkowitz, M. L.; Darden, T.; Lee, H.; Pedersen, L. G. A Smooth Particle Mesh Ewald Method. J. Chem. Phys. 1995, 103 (19), 8577–8593. https://doi.org/10.1063/1.470117.

(20) Bussi, G.; Donadio, D.; Parrinello, M. Canonical Sampling through Velocity Rescaling. J. Chem. Phys. 2007, 126 (1), 1–7. https://doi.org/10.1063/1.2408420.

(21) Parrinello, M.; Rahman, A. Polymorphic Transitions in Single Crystals : A New Molecular Dynamics Method. J. Appl. Phys. 1981, 52 (12), 7182–7190. https://doi.org/10.1063/1.328693.

(22) Hess, B. P-LINCS: A Parallel Linear Constraint Solver for Molecular Simulation. J. Chem. Theory Comput. 2008, 4 (1), 116–122. https://doi.org/10.1021/ct700200b.

(23) Michaud-agrawal, N.; Denning, E. J.; Woolf, T. B.; Beckstein, O. Software News and Updates MDAnalysis : A Toolkit for the Analysis of Molecular Dynamics Simulations. J. Comput. Chem. 2011, 32 (10), 2319–2327. https://doi.org/10.1002/jcc.

(24) Gowers, R.; Linke, M.; Barnoud, J.; Reddy, T.; Melo, M.; Seyler, S.; Domański, J.; Dotson, D.; Buchoux, S.; Kenney, I.; Beckstein, O. MDAnalysis: A Python Package for the Rapid Analysis of Molecular Dynamics Simulations. Proc. 15th Python Sci. Conf. 2016, No. Scipy, 98–105. https://doi.org/10.25080/majora-629e541a-00e.

(25) Bereau, T.; Deserno, M. Generic Coarse-Grained Model for Protein Folding and Aggregation. J. Chem. Phys. 2009, 130 (23). https://doi.org/10.1063/1.3152842.

(26) Haaga, J.; Gunton, J. D.; Buckles, C. N.; Rickman, J. M. Early Stage Aggregation of a Coarse-Grained Model of Polyglutamine. J. Chem. Phys. 2018, 148 (4). https://doi.org/10.1063/1.5010888.

(27) Bereau, T.; Globisch, C.; Deserno, M.; Peter, C. Coarse-Grained and Atomistic Simulations of the Salt-Stable Cowpea Chlorotic Mottle Virus (SS-CCMV) Subunit 26-49: β-Barrel Stability of the Hexamer and Pentamer Geometries. J. Chem. Theory Comput. 2012, 8 (10), 3750–3758. https://doi.org/10.1021/ct200888u.

(28) Bereau, T.; Bennett, W. F. D.; Pfaendtner, J.; Deserno, M.; Karttunen, M. Folding and Insertion Thermodynamics of the Transmembrane WALP Peptide. J. Chem. Phys. 2015, 143 (24). https://doi.org/10.1063/1.4935487.

(29) Rutter, G. O.; Brown, A. H.; Quigley, D.; Walsh, T. R.; Allen, M. P. Testing the Transferability of a Coarse-Grained Model to Intrinsically Disordered Proteins. Phys. Chem. Chem. Phys. 2015, 17 (47), 31741–31749. https://doi.org/10.1039/c5cp05652g.

(30) Daura, X.; Gademann, K.; Jaun, B.; Seebach, D.; Van Gunsteren, W. F.; Mark, A. E. Peptide Folding: When Simulation Meets Experiment. Angew. Chemie - Int. Ed. 1999, 38 (1-2), 236–240. https://doi.org/10.1002/(sici)1521-3773(19990115)38:1/2<236::aid-anie236>3.0.co;2-m.

(31) Adams, V. H.; McBryant, S. J.; Wade, P. A.; Woodcock, C. L.; Hansen, J. C. Intrinsic Disorder and Autonomous Domain Function in the Multifunctional Nuclear Protein, MeCP2. J. Biol. Chem. 2007, 282 (20), 15057–15064. https://doi.org/10.1074/jbc.M700855200.

(32) Kabsch, W.; Sander, C. Dictionary of Protein Secondary Structure: Pattern Recognition of Hydrogen-bonded and Geometrical Features. Biopolymers 1983, 22 (12), 2577–2637. https://doi.org/10.1002/bip.360221211.

(33) Joosten, R. P.; Te Beek, T. A. H.; Krieger, E.; Hekkelman, M. L.; Hooft, R. W. W.; Schneider, R.; Sander, C.; Vriend, G. A Series of PDB Related Databases for Everyday Needs. Nucleic Acids Res. 2011, 39 (SUPPL. 1), 411–419. https://doi.org/10.1093/nar/gkq1105.

(34) Hendrich, B.; Bird, A. Identification and Characterization of a Family of Mammalian Methyl-CpG Binding Proteins. Mol. Cell. Biol. 1998, 18 (11), 6538–6547. https://doi.org/10.1128/mcb.18.11.6538.

(35) Klose, R. J.; Sarraf, S. A.; Schmiedeberg, L.; McDermott, S. M.; Stancheva, I.; Bird, A. P. DNA Binding Selectivity of MeCP2 Due to a Requirement for A/T Sequences Adjacent to Methyl-CpG. Mol. Cell 2005, 19 (5), 667–678. https://doi.org/10.1016/j.molcel.2005.07.021.

(36) Ghosh, R. P.; Nikitina, T.; Horowitz-Scherer, R. A.; Gierasch, L. M.; Uversky, V. N.; Hite, K.; Hansen, J. C.; Woodcock, C. L. Unique Physical Properties and Interactions of the Domains of Methylated DNA Binding Protein 2. Biochemistry 2010, 49 (20), 4395–4410. https://doi.org/10.1021/bi9019753.

(37) Ghosh, R. P.; Horowitz-Scherer, R. A.; Nikitina, T.; Gierasch, L. M.; Woodcock, C. L. Rett Syndrome-Causing Mutations in Human MeCP2 Result in Diverse Structural Changes That Impact Folding and DNA Interactions. J. Biol. Chem. 2008, 283 (29), 20523–20534. https://doi.org/10.1074/jbc.M803021200.

(38) Heinig, M.; Frishman, D. STRIDE: A Web Server for Secondary Structure Assignment from Known Atomic Coordinates of Proteins. Nucleic Acids Res. 2004, 32 (WEB SERVER ISS.), 500–502. https://doi.org/10.1093/nar/gkh429.

(39) Kim, H.; Park, H. Prediction of Protein Relative Solvent Accessibility with Support Vector Machines and Long-Range Interaction 3D Local Descriptor. Proteins Struct. Funct. Genet. 2004, 54 (3), 557–562. https://doi.org/10.1002/prot.10602.

(40) Gress, A.; Kalinina, O. V. SphereCon - A Method for Precise Estimation of Residue Relative Solvent Accessible Area from Limited Structural Information. Bioinformatics 2020, 36 (11), 3372–3378. https://doi.org/10.1093/bioinformatics/btaa159.

(41) Ho, K. L.; McNae, I. W.; Schmiedeberg, L.; Klose, R. J.; Bird, A. P.; Walkinshaw, M. D. MeCP2 Binding to DNA Depends upon Hydration at Methyl-CpG. Mol. Cell 2008, 29 (4), 525–531. https://doi.org/10.1016/j.molcel.2007.12.028.

(42) Lei, M.; Tempel, W.; Chen, S.; Liu, K.; Min, J. Plasticity at the DNA Recognition Site of the MeCP2 MCG-Binding Domain. Biochim. Biophys. Acta - Gene Regul. Mech. 2019, 1862 (9), 194409. https://doi.org/10.1016/j.bbagrm.2019.194409.

(43) Humphrey, W.; Dalke, A.; Schulten, K. VMD: Visual Molecular Dynamics. J. Mol. Graph. 1996, 14 (1), 33–38. https://doi.org/10.1016/0263-7855(96)00018-5.

(44) Schrödinger, L. The PyMOL Molecular Graphics System, Version 2.0 Schrödinger, LLC. 2015.

(45) Bonomi, M.; Branduardi, D.; Bussi, G.; Camilloni, C.; Provasi, D.; Raiteri, P.; Donadio, D.; Marinelli, F.; Pietrucci, F.; Broglia, R. A.; Parrinello, M. PLUMED: A Portable Plugin for Free-Energy Calculations with Molecular Dynamics. Comput. Phys. Commun. 2009, 180 (10), 1961–1972. https://doi.org/10.1016/j.cpc.2009.05.011.

(46) Pietrucci, F.; Laio, A. A Collective Variable for the Efficient Exploration of Protein Beta-Sheet Structures: Application to SH3 and GB1. J. Chem. Theory Comput. 2009, 5 (9), 2197–2201. https://doi.org/10.1021/ct900202f.

(47) Šolc, K. Shape of a Random-Flight Chain. J. Chem. Phys. 1971, 55 (1).

(48) Tolmachev, D. A.; Boyko, O. S.; Lukasheva, N. V.; Martinez-Seara, H.; Karttunen, M. Overbinding and Qualitative and Quantitative Changes Caused by Simple Na+ and K+ Ions in Polyelectrolyte Simulations: Comparison of Force Fields with and without NBFIX and ECC Corrections. J. Chem. Theory Comput. 2020, 16 (1), 677–687. https://doi.org/10.1021/acs.jctc.9b00813.

(49) Rauscher, S.; Gapsys, V.; Gajda, M. J.; Zweckstetter, M.; De Groot, B. L.; Grubmüller, H. Structural Ensembles of Intrinsically Disordered Proteins Depend Strongly on Force Field: A Comparison to Experiment. J. Chem. Theory Comput. 2015, 11 (11), 5513–5524. https://doi.org/10.1021/acs.jctc.5b00736.

(50) Chang, M.; Wilson, C. J.; Karunatilleke, N. C.; Moselhy, M. H.; Karttunen, M.; Choy, W. Y. Exploring the Conformational Landscape of the Neh4 and Neh5 Domains of Nrf2 Using Two Different Force Fields and Circular Dichroism. J. Chem. Theory Comput. 2021, 17 (5), 3145–3156. https://doi.org/10.1021/acs.jctc.0c01243.

(51) Tunyasuvunakool, K.; Adler, J.; Wu, Z.; Green, T.; Zielinski, M.; Žídek, A.; Bridgland, A.; Cowie, A.; Meyer, C.; Laydon, A.; Velankar, S.; Kleywegt, G. J.; Bateman, A.; Evans, R.; Pritzel, A.; Figurnov, M.; Ronneberger, O.; Bates, R.; Kohl, S. A. A.; Potapenko, A.; Ballard, A. J.; Romera-Paredes, B.; Nikolov, S.; Jain, R.; Clancy, E.; Reiman, D.; Petersen, S.; Senior, A. W.; Kavukcuoglu, K.; Birney, E.; Kohli, P.; Jumper, J.; Hassabis, D. Highly Accurate Protein Structure Prediction for the Human Proteome. Nature 2021, 596 (7873), 590–596. https:x//doi.org/10.1038/s41586-021-03828-1.

(52) Ruff, K. M.; Pappu, R. V. AlphaFold and Implications for Intrinsically Disordered Proteins. J. Mol. Biol. 2021, 433 (20), 167208. https://doi.org/10.1016/j.jmb.2021.167208.

(53) Wilson, C. J.; Choy, W.; Karttunen, M. AlphaFold2 : A Role for Disordered Protein Prediction? bioRxiv 2021. https://doi.org/https://doi.org/10.1101/2021.09.27.461910.

(54) Strodel, B. Energy Landscapes of Protein Aggregation and Conformation Switching in Intrinsically Disordered Proteins. J. Mol. Biol. 2021, No. xxxx. https://doi.org/10.1016/j.jmb.2021.167182.

(55) Kodera, N.; Noshiro, D.; Dora, S. K.; Mori, T.; Habchi, J.; Blocquel, D.; Gruet, A.; Dosnon, M.; Salladini, E.; Bignon, C.; Fujioka, Y.; Oda, T.; Noda, N. N.; Sato, M.; Lotti, M.; Mizuguchi, M.; Longhi, S.; Ando, T. Structural and Dynamics Analysis of Intrinsically Disordered Proteins by High-Speed Atomic Force Microscopy. Nat. Nanotechnol. 2021, 16 (2), 181–189. https://doi.org/10.1038/s41565-020-00798-9.

(56) Solanki, A.; Neupane, K.; Woodside, M. T. Single-Molecule Force Spectroscopy of Rapidly Fluctuating, Marginally Stable Structures in the Intrinsically Disordered Protein α - Synuclein. Phys. Rev. Lett. 2014, 112 (15), 1–6. https://doi.org/10.1103/PhysRevLett.112.158103.

(57) Cheng, S.; Cetinkaya, M.; Gräter, F. How Sequence Determines Elasticity of Disordered Proteins. Biophys. J. 2010, 99 (12), 3863–3869. https://doi.org/10.1016/j.bpj.2010.10.011.

(58) Ando, T.; Uchihashi, T.; Scheuring, S. Filming Biomolecular Processes by High-Speed Atomic Force Microscopy. Chem. Rev. 2014, 114 (6), 3120–3188. https://doi.org/10.1021/cr4003837.

