## Supplementary Information for "A multiscale computational study of the conformation of the full-length intrinsically disordered protein MeCP2"

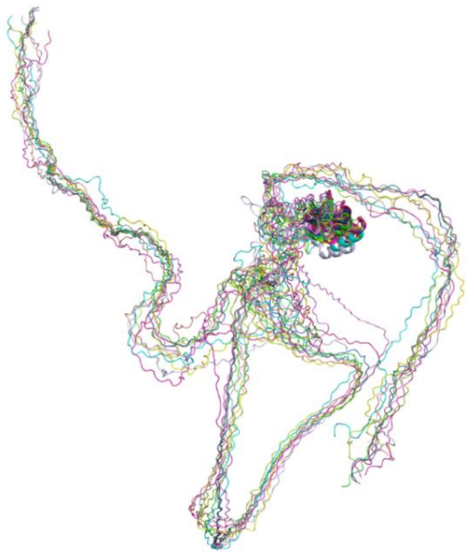

Figure S1. Alignment of ten of the rejected models of the full-length MeCP2 protein built with Modeller[18].

Table S1. Details of all-atom simulations.

| *System* | *Water molecules* | *Na/Cl each* | *Counterions (Cl)* | *Duration (ns)* | *Starting structure* |
|---|---|---|---|---|---|
| *MeCP2_1* | 793,027 | 2,205 | 37 | 150 | Built with Modeller |
| *MeCP2_1 resized* | 119,575 | 337 | 37 | 1000 | MeCP2_1 |
| *MeCP2_2* | 113,980 | 324 | 37 | 60 | Built with Modeller |
| *MeCP2_3* | 458,344 | 1,277 | 37 | 60 | Built with Modeller |
| *MeCP2_3 resized* | 119,989 | 339 | 37 | 600 | MeCP2_3 |
| *Loop Amber* | 324,046 | 0 | 31 | 10 | loop in MeCP2_3 |
| *Loop A1 resized* | 71,863 | 0 | 31 | 100 | Loop Amber |
| *Loop A2 resized* | 44,741 | 0 | 31 | 100 | Loop Amber |
| *Loop A3 resized* | 224,502 | 0 | 31 | 100 | Loop Amber |

| | | | | | |
|---|---|---|---|---|---|
| *Loop A4 resized* | 129,396 | 0 | 31 | 100 | Loop Amber |
| *Loop A5 resized* | 131,425 | 0 | 31 | 100 | Loop Amber |
| *Loop Charmm* | 324,046 | 0 | 31 | 10 | loop in MeCP2_3 |
| *Loop C1 resized* | 186,560 | 0 | 31 | 100 | Loop Charmm |
| *Loop C2 resized* | 281,911 | 0 | 31 | 100 | Loop Charmm |
| *Loop C3 resized* | 54,506 | 0 | 31 | 100 | Loop Charmm |
| *Loop C4 resized* | 293,861 | 0 | 31 | 100 | Loop Charmm |
| *Loop C5 resized* | 102,059 | 0 | 31 | 100 | Loop Charmm |
| *Loop 228-242* | 4,665 | 0 | 0 | 100 | Loop in two-globule, CG1 |
| *Loop 161-205* | 13,658 | 0 | 12 | 100 | Loop in two-globule, CG4 |
| *AlphaFold R1* | 144,569 | 411 | 37 | 400 | AlphaFold model |
| *AlphaFold R2* | 144,569 | 411 | 37 | 400 | AlphaFold model |
| *AlphaFold R3* | 144,569 | 411 | 37 | 400 | AlphaFold model |

Table S2. Details of coarse-grained simulations. These were run using the PLUM model with implicit water.

| *System* | *Duration (ns)* |
|---|---|
| *CG1* | 3,000 |
| *CG2* | 500 |

| | |
|---|---|
| *CG3* | 500 |
| *CG4* | 700 |

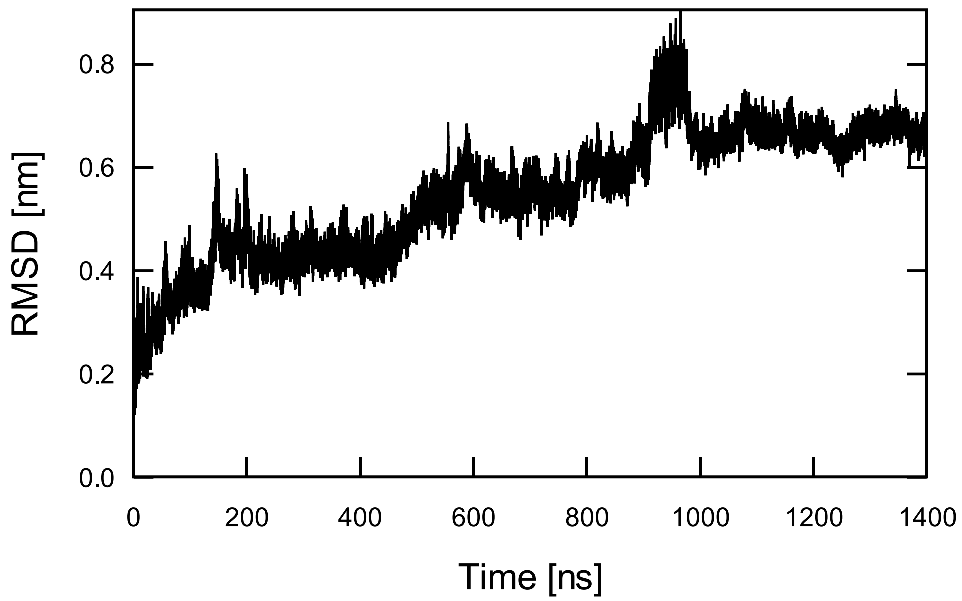


Figure S2. RMSD of the TRD domain in the all-atom full-length protein simulation.

Table S3. Secondary structure content in the last 400 ns of the all-atom MeCP2_1 simulation clustered with a 0.5 nm cutoff and the Daura *et al.* method[39].

| *Cluster #* | *# frames* | *α-helix* | *β-strand/turn* | *coil/bend* |
|---|---|---|---|---|
| *1* | 23,559 | 22 | 121 | 333 |
| *2* | 11,556 | 16 | 99 | 353 |
| *3* | 2,920 | 26 | 110 | 338 |
| *4* | 1,189 | 15 | 112 | 342 |
| *5* | 499 | 31 | 109 | 327 |
| *6* | 137 | 8 | 120 | 340 |
| *7* | 58 | 14 | 109 | 343 |

| | | | | |
|---|---|---|---|---|
| *8* | 58 | 21 | 109 | 338 |
| *9* | 19 | 23 | 101 | 345 |
| *10* | 3 | 15 | 115 | 343 |
| *11* | 3 | 22 | 91 | 350 |
| *Total/Average* | 40,001 | 20 | 113 | 339 |

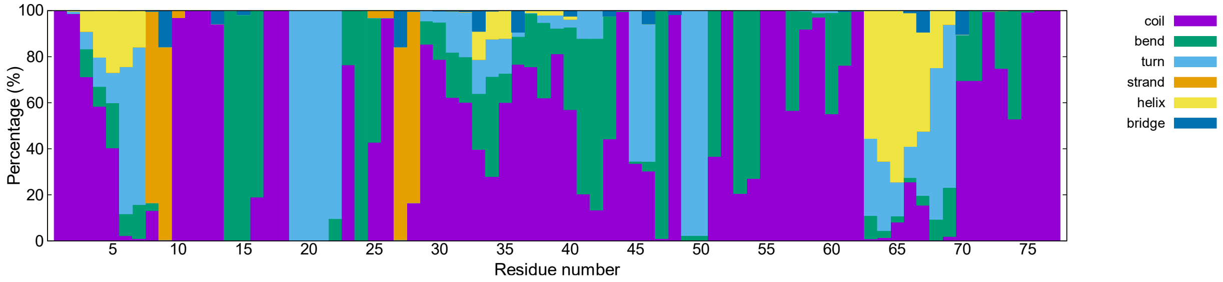


Figure S3. Percentage of frames in the last 400 ns of the NTD domain with every type of secondary structure.

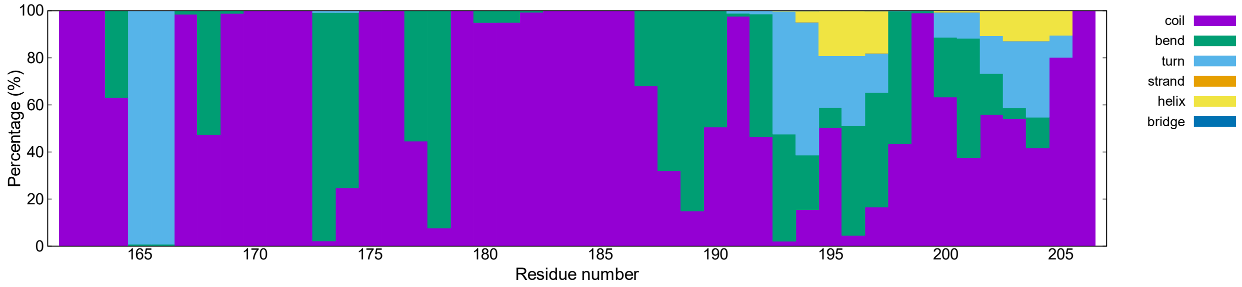


Figure S4. Percentage of frames in the last 400 ns of the ID domain with every type of secondary structure.

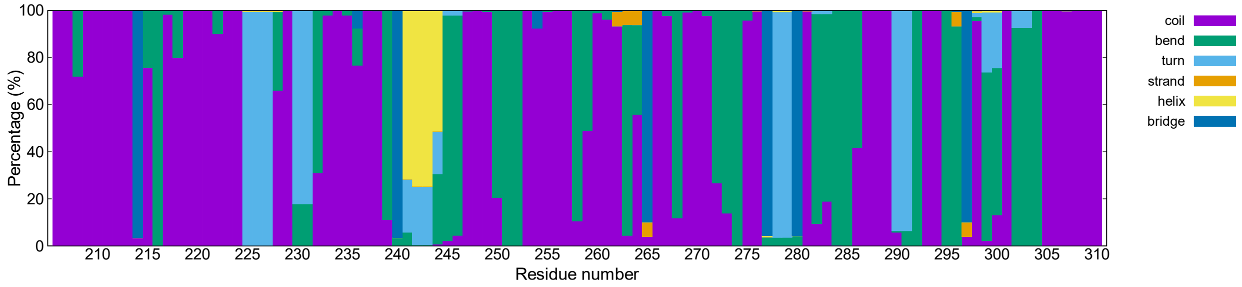

Figure S5. Percentage of frames in the last 400 ns of the TRD domain with every type of secondary structure.

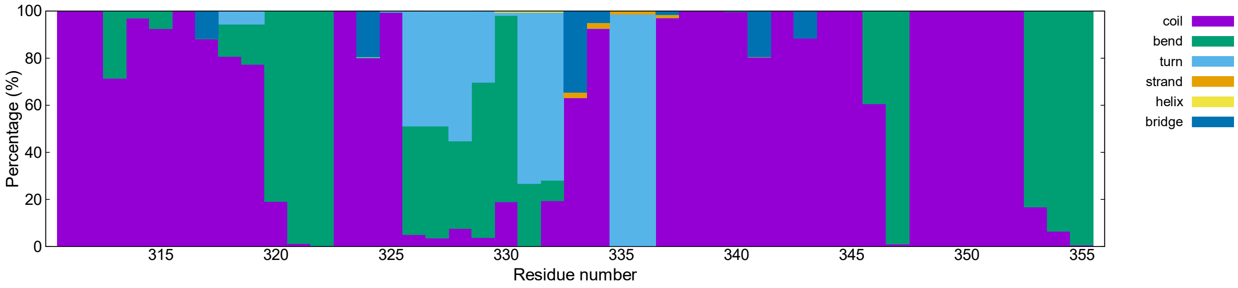


Figure S6. Percentage of frames in the last 400 ns of the CTDα domain with every type of secondary structure.

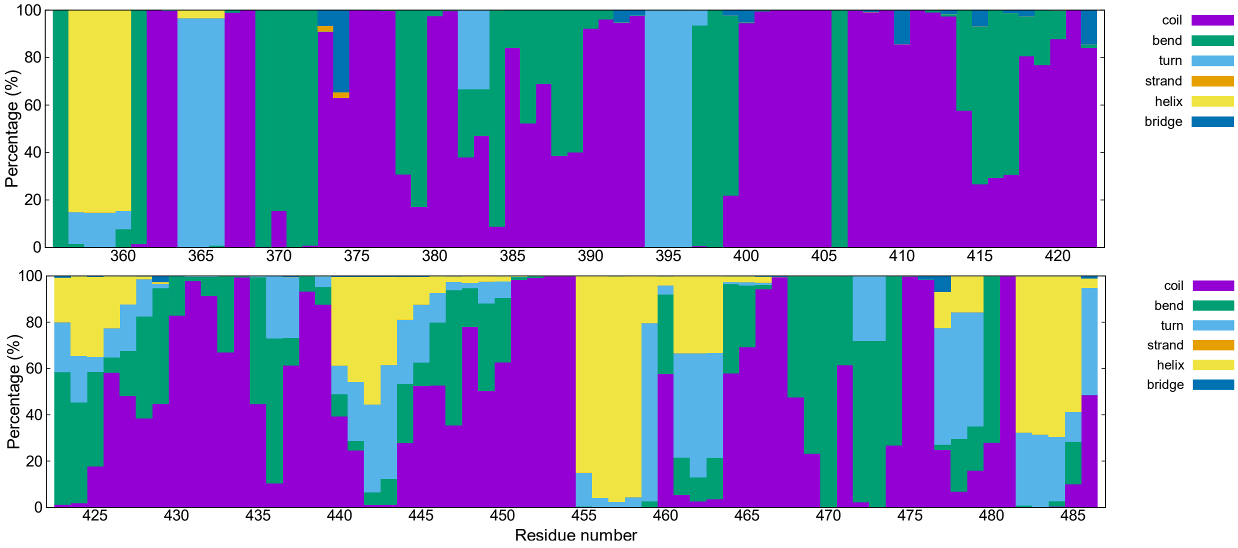


Figure S7. Percentage of frames in the last 400 ns of the CTDβ domain with every type of secondary structure.

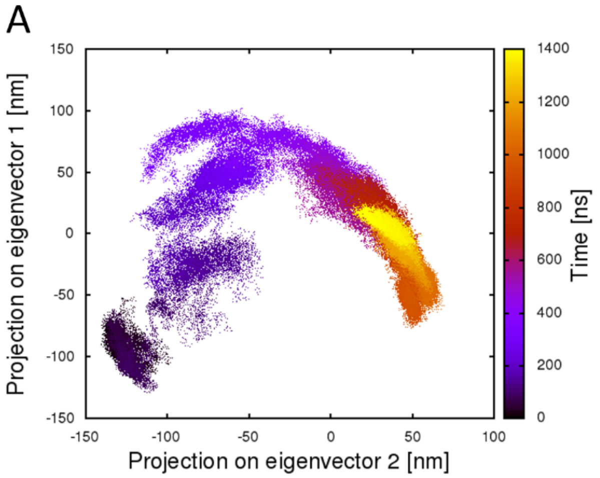


Figure S8. Principal Component Analysis. Projection of the MeCP2_1 simulation on the first two eigenvectors, coloured by time. (A) Projection of the entire protein. (B) Projection of the TRD domain. The protein samples the largest portion of this 2D-space during the first 600 ns of the simulation. In contrast, the TRD domain begins to sample more conformational space in the second half of the simulation.

Table S4. Relative solvent accessible surface area (rSA) of residue W104 in the four most populated clusters of the all-atom simulation.

| *Cluster #* | *weight* | *rASA* |
|---|---|---|
| *1* | 23,559 | 7.4 |
| *2* | 11,556 | 9.6 |
| *3* | 2,920 | 6.6 |
| *4* | 1,189 | 11.4 |
| | Weighted average: | 8.1 |

Table S5. Relative solvent accessible surface area (rSA) of residue R111 in the four most populated clusters of the all-atom simulation.

| *Cluster #* | *weight* | *rASA* |
|---|---|---|
| *1* | 23,559 | 12.7 |
| *2* | 11,556 | 37.4 |

| | | |
|---|---|---|
| *3* | 2,920 | 5.8 |
| *4* | 1,189 | 44.3 |
| | Weighted average: | 20.4 |

Table S6. Relative solvent accessible surface area (rSA) of residue R133 in the four most populated clusters of the all-atom simulation.

| *Cluster #* | *weight* | *rASA* |
|---|---|---|
| *1* | 23,559 | 50.3 |
| *2* | 11,556 | 56.7 |
| *3* | 2,920 | 65.5 |
| *4* | 1,189 | 63.0 |
| | Weighted average: | 53.7 |

Table S7. Salt bridges in the last 400 ns of the full-length all-atom MeCP2_1 simulation.

| Interacting residues | Protein domains |
|---|---|
| *Glu11 – Lys27* | NTD – NTD |
| *Asp15 – Arg162* | NTD – MBD |
| *Asp17 – Lys135* | NTD – MBD |
| *Lys22 – Glu214* | NTD – TRD |
| *Glu55 – Lys109* | NTD – MBD |
| *Glu55 – Arg111* | NTD – MBD |
| *Glu55 – Arg133* | NTD – MBD |
| *Glu66 – Arg91* | NTD – MBD |

| | |
|---|---|
| *Glu66 – Arg85* | NTD – MBD |
| *Glu76 – Arg85* | NTD – MBD |
| *Asp97 – Lys171* | MBD – ID |
| *Arg111 – Asp121* | MBD – MBD |
| *Asp154 – Arg167* | MBD – ID |
| *Asp156 – Arg167* | MBD – ID |
| *Arg168 – Glu205* | ID – ID |
| *Glu235 – Lys254* | TRD – TRD |
| *Glu235 – Lys347* | TRD – CTDα |
| *Arg253 – Glu315* | TRD – CTDα |
| *Arg253 – Glu365* | TRD – CTDβ |
| *Arg255 – Glu258* | TRD – TRD |
| *Asp260 – Lys337* | TRD – CTDα |
| *Lys266 – Glu282* | TRD – TRD |
| *Arg270 – Glu394* | TRD – CTDβ |
| *Lys271 – Glu404* | TRD – CTDβ |
| *Glu290 – Lys307* | TRD – TRD |
| *Arg294 – Glu318* | TRD – CTDα |
| *Glu298 – Lys307* | TRD – TRD |
| *Lys304 – Asp407* | TRD – CTDβ |
| *Lys321 – Glu397* | CTDα – CTDβ |
| *Arg344 – Glu365* | CTDα – CTDβ |
| *Arg420 – Glu473* | CTDβ – CTDβ |
| *Glu432 – Arg453* | CTDβ – CTDβ |

| | |
|---|---|
| *Arg453 – Glu457* | CTDβ – CTDβ |
| *Glu455 – Arg458* | CTDβ – CTDβ |
| *Glu457 – Arg471* | CTDβ – CTDβ |

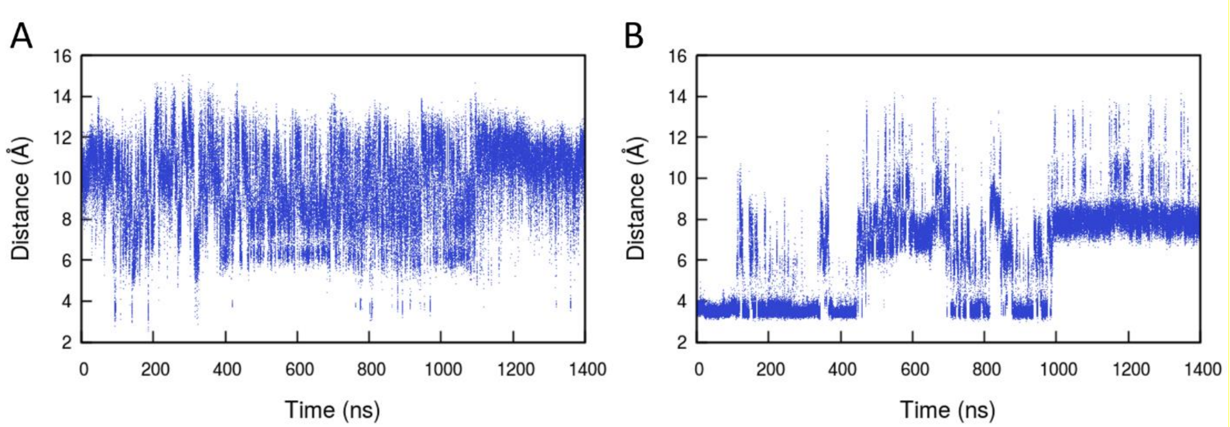


Figure S9. Salt bridge interaction between residues Lys 119 and Asp 121 (A) and between residues Arg 133 and Glu 137 (B).

Table S8. Clusters sampled in the coarse-grained simulations. The first eight clusters contain 95.3% of the sampled structures.

| *Cluster #* | *# Structures* | *Sampled in* | *Representative structure* |
|---|---|---|---|
| *1* | 22,293 | CG1, CG2, CG4 | 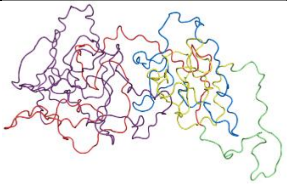 |
| *2* | 5,810 | CG1, CG2, CG3, CG4 | 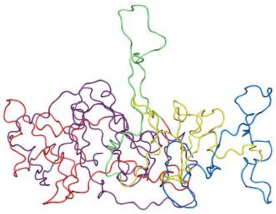 |

| | | | |
|---|---|---|---|
| *3* | 4,986 | CG1 | 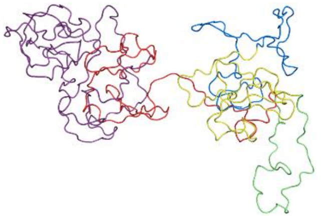 |
| *4* | 3,492 | CG1, CG3 | 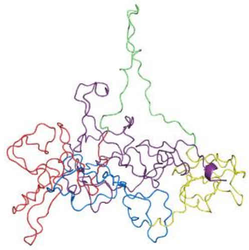 |
| *5* | 3,027 | CG1, CG4 | 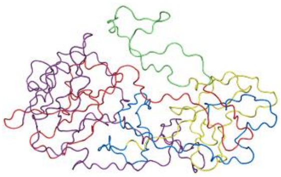 |
| *6* | 2,768 | CG4 | 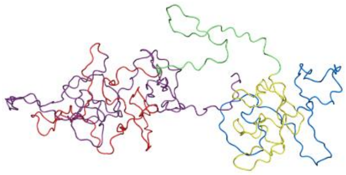 |
| *7* | 1,257 | CG4 | 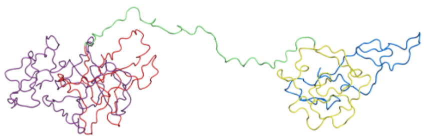 |
| *8* | 1,165 | CG1 | 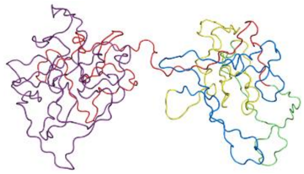 |

Table S9. Templates used to generate models MeCP2_2 and MeCP2_3. The blue templates were taken from the coarse-grained simulation CG1, the green template was generated with Pymol[52], the red template was taken from an all-atom simulation of the NTD+MBD domains[66] and the purple template was taken from model MeCP2_2.

| | **TEMPLATE 1** | **TEMPLATE 2** | **TEMPLATE 3** | **FINAL MODEL** |
|---|---|---|---|---|
| **MECP2_2** | 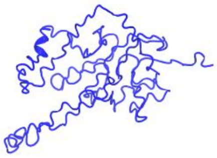 | 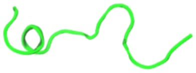 | 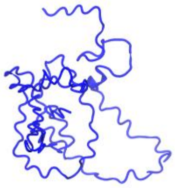 | 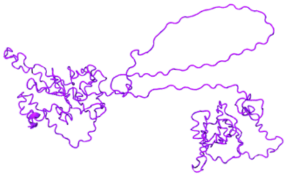 |
| **MECP2_3** | 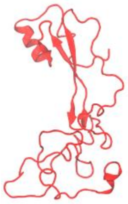 | 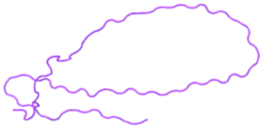 | 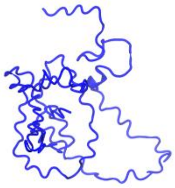 | 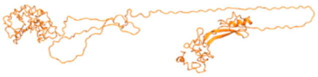 |

Table S10. Conformations sampled by the ID+TRD domains simulations. The five most populated clusters are shown for each simulation.

| | **Replica 1** | **Replica 2** | **Replica 3** | **Replica 4** | **Replica 5** |
|---|---|---|---|---|---|
| **Amber** | 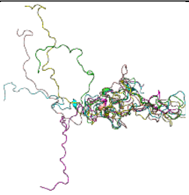 | 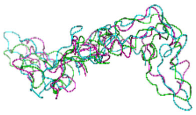 | 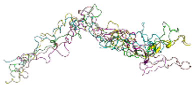 | 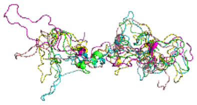 | 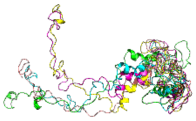 |
| **Charmm** | 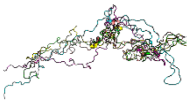 | 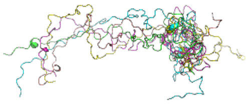 | 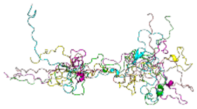 | 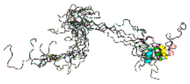 | 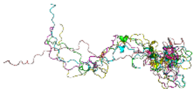 |

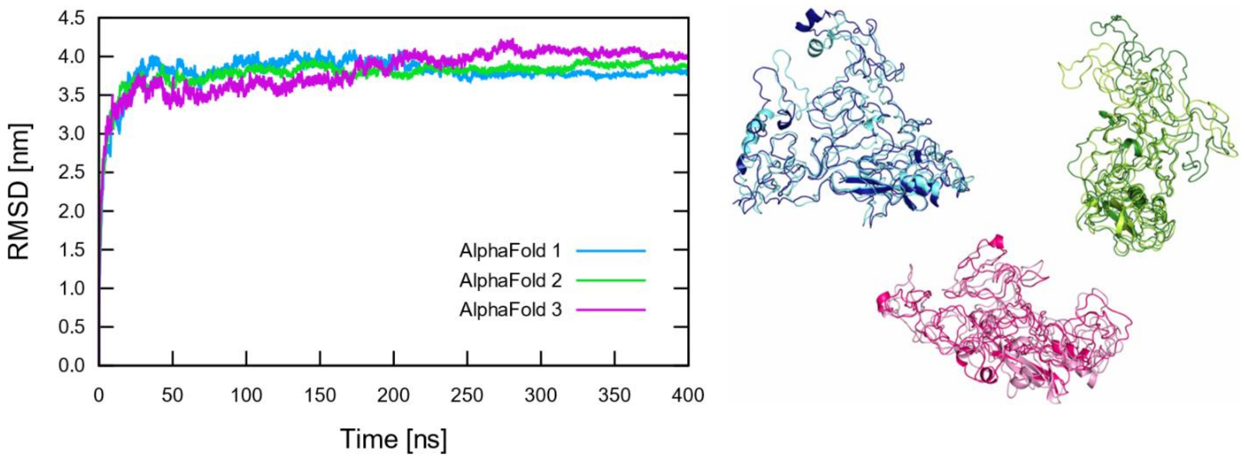


Figure S10. RMSD of the three AlphaFold simulations. Right: Alignment of the most sampled structure from 200 to 300 ns (light colours) and the most sampled structure in the last 100 ns (dark colours) of each replica. Both structures are very similar to each other in the three simulations, ratifying the convergence of the simulations.
